# Manipulation-specific activity in motor and somatosensory cortex as mice handle food

**DOI:** 10.1101/2022.02.23.481687

**Authors:** John M. Barrett, Gordon M. G. Shepherd

## Abstract

Food-handling offers unique yet largely unexplored opportunities to investigate how cortical activity relates to forelimb movements in a natural, ethologically essential, and kinematically rich form of manual dexterity. To determine these relationships, we recorded spiking activity in mouse forelimb M1 and S1 and tongue/jaw M1. Activity in all areas was strongly modulated in close association with discrete active manipulation events that occurred intermittently as mice fed. Each area’s activity was also partly distinct in its overall timing and phasic/tonic temporal profile, attributable to area-specific composition of activity classes. Forelimb position could be accurately predicted from activity in all three regions. These results thus establish that cortical activity during food-handling is manipulation-specific, distributed, and broadly similar across multiple cortical areas, while also exhibiting area- and submovement-specific relationships with the fast kinematic hallmarks of this form of complex, free-object-handling manual dexterity.

## INTRODUCTION

Manual dexterity takes many forms (Sobinov and Bensmaia, 2021). One, prominent in primates and rodents including mice, is food-handling (Whishaw and Coles, 1996; Barrett et al., 2020). Food-handling for these species is ethologically critical, constituting a basic and essential form of manual dexterity. Food-handling entails tight interaction between motor output and sensory input as the morsel is manipulated and consumed. Food-handling is kinematically rich, with rapid coordinated movements of the forelimbs and orofacial structures. These properties make food-handling an attractive behavior for studying the neurobiology of manual dexterity. For this, mice hold promise as experimentally tractable model organisms. Mice handle food similarly whether head-fixed or freely moving, and the basic kinematic features of food-handling are characterized (Barrett et al., 2020). For other forms of manual dexterity in mice, such as reach-to-grasp and learned manipulandum-based tasks, knowledge is rapidly advancing about the associated neural circuits and systems (Warriner et al., 2020). Characterizing these for mouse food-handling could provide fundamental insights into the neurobiology of not only this behavior but manual dexterity in general.

Forelimb motor cortex (M1) is involved in many forms of manual dexterity and thus presents a starting point for investigating cortical roles in food-handling. On the one hand, M1 involvement might be minimal: motor cortex disengages over time on well-learned tasks (Hwang et al., 2019; Hwang et al., 2021), and brainstem stimulation alone can evoke food-handling-like movements (Ruder et al., 2021). On the other hand, cortical stimulation in motor areas also evokes food-handling-like behaviors in primates (Graziano et al., 2002) and mice (Hira et al., 2015; Mercer Lindsay et al., 2019); motor cortical lesions can impair food-handling in rats (Whishaw and Coles, 1996); and, M1 neurons exhibit diverse movement-related activity (Fromm and Evarts, 1977; Miri et al., 2017; Sjöbom et al., 2020; Guo et al., 2021). M1 activity associated with food-handling movements could take many forms. At one extreme, individual neurons could be heterogeneously active, averaging out at the population level. At another, neuronal activity could be homogeneous, fluctuating as a population. A third possibility is a hybrid pattern, combining elements of neuronal heterogeneity and population-wide fluctuations. Additionally, other cortical motor and somatosensory areas are likely involved, particularly forelimb S1 and tongue/jaw M1 (Mayrhofer et al., 2019), and might be differentiated in terms of their precise timing and temporal profiles. Given the high speed of food-handling movements (Barrett et al., 2020), distinguishing among these hypothetical possibilities will require recordings of neural activity and kinematics with high temporal resolution.

Here, we used multielectrode array electrophysiological recordings from forelimb M1 and high-speed videography to capture neuronal spiking and kinematics at high time resolution while mice handled food. We also extended the analysis to forelimb S1 and tongue/jaw M1. To make sense of the large, complex datasets, we used analytical methods that enabled assessment of both neuron- and population-level aspects of the activity patterns. Our results establish the basic properties of food-handling related cortical activity in motor and somatosensory areas, showing robust modulation of activity in multiple areas specifically during active manipulation events, with distinct temporal profiles for each region.

## RESULTS

### Forelimb M1 activity during food-handling is associated with oromanual events

To investigate cortical activity during food-handling, we presented head-fixed mice with food items (sunflower seed kernels or grain pellets) and filmed their movements with close-up dual-angle kilohertz-rate video while recording spiking activity from all cortical layers on multi-channel linear electrode arrays (one shank per array, 32 channels per shank, 50 µm spacing) as they handled and consumed the morsels (**Fig. 1, Videos 1-2, Methods**). To capture kinematic and cortical activity on the extended time scale of sustained food-handling, recordings were made over tens of seconds (range: 11.7 to 97.5 sec).

**Fig. 1:**
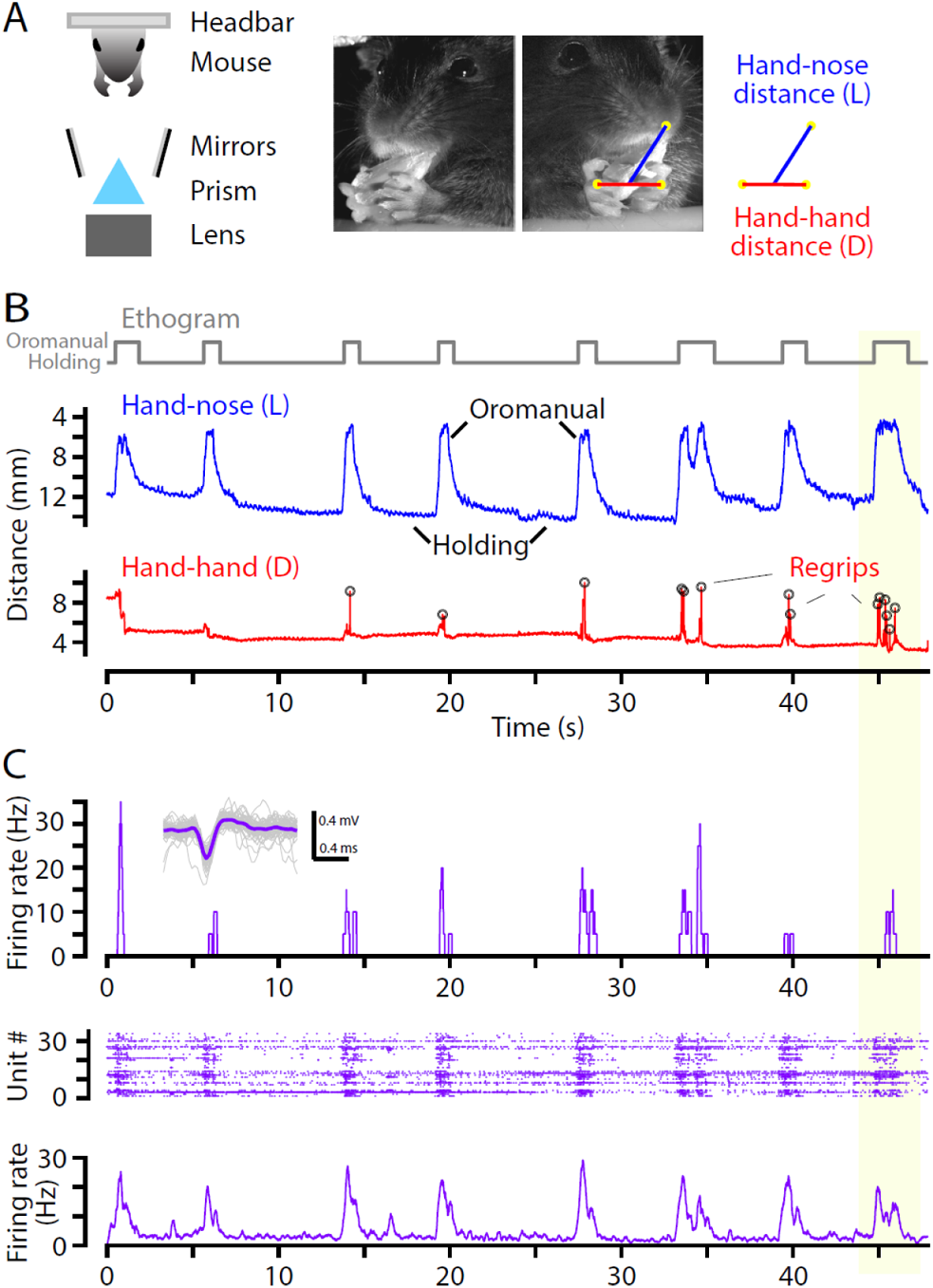
Forelimb M1 activity during food-handling is associated with oromanual events. (A) Kinematics were recorded by a close-up kilohertz-rate video system, and cortical activity was recorded on multi-channel linear arrays, while head-fixed mice handled food items (left). Video frames (middle) were analyzed to track hand movements, and kinematic data were analyzed to calculate the hand-nose (L) and hand-hand (D) distances (right). (B) Example trace of the hand-nose distance (L, blue, reverse y-axis) over time, showing characteristic “oromanual” events (intermittent peaks in L), separated by a “holding” postural mode (large L). The trace of the hand-hand distance (D, red) shows fast spikes corresponding to “regrip” events. Ethogram (top) shows oromanual events and holding modes. Yellow boxed region (right) highlights one example of an oromanual event. (C) Traces of corresponding electrophysiological activity, showing bursts of firing associated with oromanual events. Top: plot of firing rate for a single unit recorded on one channel. Inset: Example single unit. Middle: raster plot of spiking recorded on multiple channels on the same probe. Bottom: average firing rate across all units recorded on the same probe.

We analyzed the videos using DeepLabCut (Mathis et al., 2018) to track movements of the hands and nose, and reconstructed their trajectories in 3D using Anipose (Karashchuk et al., 2020), from which we extracted kinematic parameters of interest. As previously (Barrett et al., 2020), we focused on the three-dimensional Euclidean hand-nose distance (L), and the 3D hand-hand distance (D) (**Fig. 1A**). Plotting these parameters over time (with L on a reverse Y axis so that upward movements of the hands correspond to upward movements of the trace) showed several characteristic features (**Fig. 1B**). These included frequent broad peaks in L indicating “oromanual” events in which the hands brought the food item to the mouth, longer “holding” intervals in which the food item was held passively below the mouth, and intermittent fast spikes in D, observed only during oromanual events, as the hands quickly readjusted their grip on the food (“regrips”).

Electrophysiological recordings from multi-channel linear arrays placed in forelimb M1 were analyzed to extract single- and multi-unit spiking activity (**Fig. 1C, Methods**). From each recording we isolated 14 ± 12 (mean ± s.d.) single- and 32 ± 9 multi-units per array. Single- and multi-units (“active units”) were included in all analyses of spiking activity. Plotting the firing rate over time showed frequent bursts of activity, aligned with oromanual events. Such patterns were apparent across recordings (**Fig. S1**). Accordingly, we created oromanual/holding ethograms from the L traces and from these calculated the average firing rates during these periods. Across animals, the firing rate was twice as high during oromanual events compared to holding (holding 5.5 ± 2.0 Hz, mean ± s.d.; oromanual 9.9 ± 4.5 Hz; *W* = 0, *p* = 0.031, Wilcoxon signed-rank test, *n* = 6 mice). However, averaging across such large temporal windows obscures the fine details of the relationship between cortical activity and food-handling kinematics. We therefore next analyzed the kinematic features of oromanual events in finer detail.

### Kinematic composition of oromanual events

Closer inspection of oromanual events shows they are composed of several distinct kinematic features (**Fig. 2A**). These include a rapid, upward, “transport-to-mouth” movement (reduction in L) as the food is brought to the mouth, and a slower, downward, “lowering-from-mouth” movement (increase in L) as the hands drop to a holding posture while the mouse chews. Regrips (spikes in D) occur during oromanual events.

**Fig. 2:**
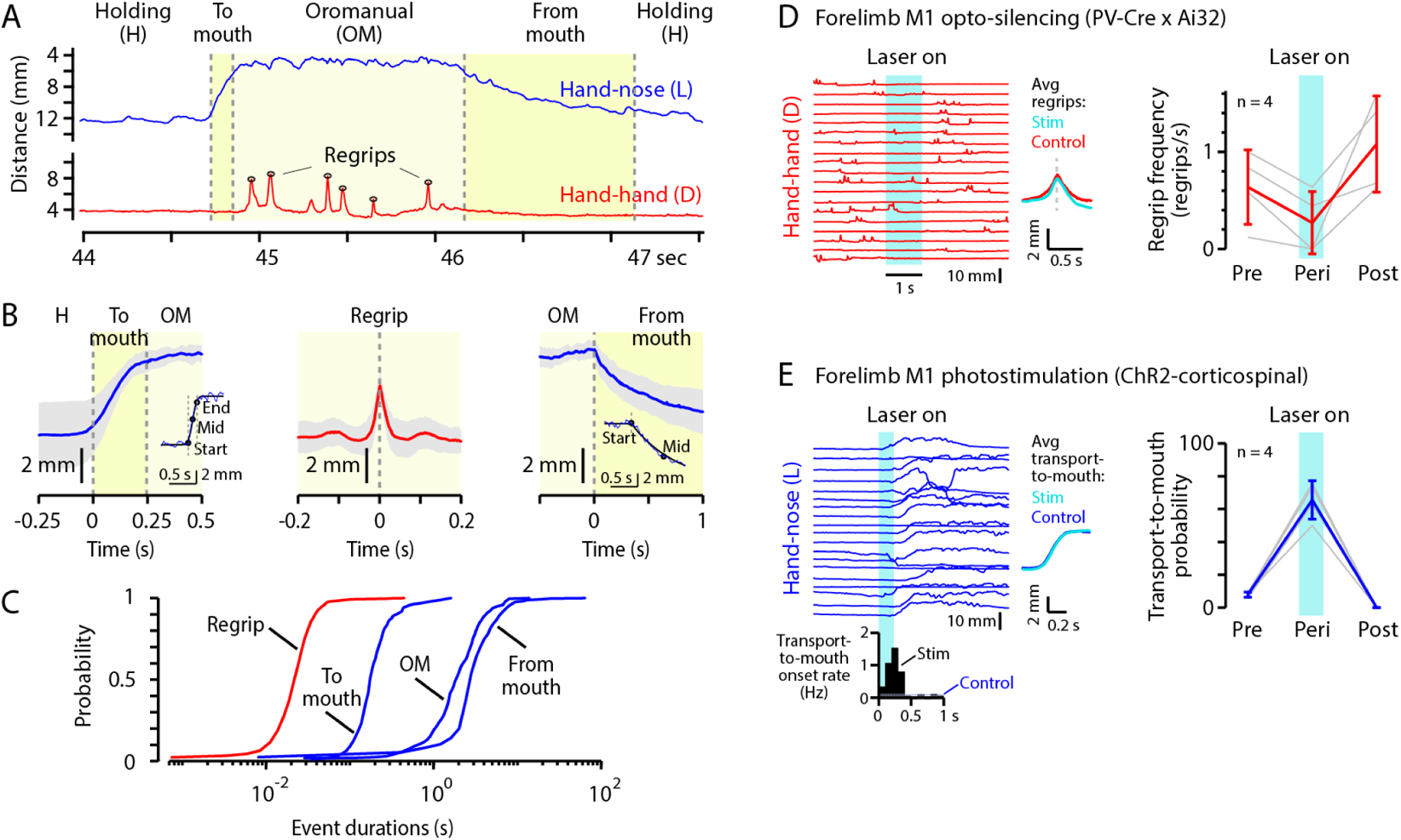
Kinematic composition of oromanual events. (A) Example of an oromanual event (last event in the example shown in Fig. 1), characterized by a rapid transport-to-mouth movement, multiple regrips during the oromanual period, and a slower lowering-from-mouth movement. (B) Plots show event-alignment and averaging of different features of the oromanual events. Left: Average transport-to-mouth movement, aligned to the onset of the change in L. Error bands indicate mean ± s.d for *n* = 9 mice. Inset shows an example of fitting a curve to identify the start, middle, and end points of the transport-to-mouth movement (**Methods**). Middle: Average regrip, aligned to the peak of the spike in D. Right: Average lowering-from-mouth movement, aligned to the onset of the change in L. Inset shows an example of curve-fitting to identify the key time points of the lowering-from-mouth movement. (C) Cumulative probability plots of the durations of regrips, transport-to-mouth movements, oromanual durations, and lowering-from-mouth movements. Note semi-log scale. (D) Left: Example D traces from an optogenetic silencing experiment. Blue bar represents the duration of the laser stimulus. The two traces to the right show the average regrip trajectories between silencing trials (red, *n* = 4 mice) and during silencing trials (cyan, *n* = 2 mice with regrips during silencing). Right: Frequency of regrips during oromanual events in the 2 seconds before (‘Pre’), 1 second during (‘Peri’), and 2 seconds after (‘Post’) silencing. Thin grey lines are individual mice (*n* = 4) and thick red error bars are mean ± s.d. over mice. (E) Left: Example L traces from an optogenetic stimulation experiment. Blue bar represents the duration of the laser stimulus. The peristimulus time histogram below shows the rate of transport-to-mouth movements in 100 ms bins following stimulus onset when stimulation was delivered during holding for *n* = 4 mice, with the background rate (‘Control’) indicated by thin horizontal lines (blue dashed line: mean, solid grey lines: s.d.). The two traces to the right show the average transport-to-mouth trajectories immediately following stimulation (cyan) and at all other times (blue). Right: probability of observing a transport-to-mouth movement within 400 ms of stimulus onset (‘Peri’) compared to virtual stimulus timings shifted one second earlier (‘Pre’) or later (‘Post’). Only trials where the stimulus (real or virtual) occurred during holding are included. Thin grey lines are individual mice (*n* = 4) and thick blue error bars are mean ± s.d. over mice.

To quantify these features, we first aligned to threshold crossings in L. This showed a sigmoid-like trajectory for the transport-to-mouth movement and an exponential-like trajectory for the lowering-from-mouth movement. Accordingly, we fit each transition movement with the corresponding function and used these to quantify the timing (**Methods**). For the hand-nose distance, L, alignment to the onset of the transport-to-mouth movement showed an average amplitude of 4.7 ± 1.7 mm (mean ± s.d., *n* = 9 mice, **Table 1**) and duration of 246 ± 116 ms (**Fig. 2B, left**). At the end of oromanual events, the lowering-from-mouth movement had a similar average amplitude of 4.6 ± 1.5 mm but a longer duration of 4.0 ± 1.1 s (**Fig. 2B, right**).

**Table 1:**
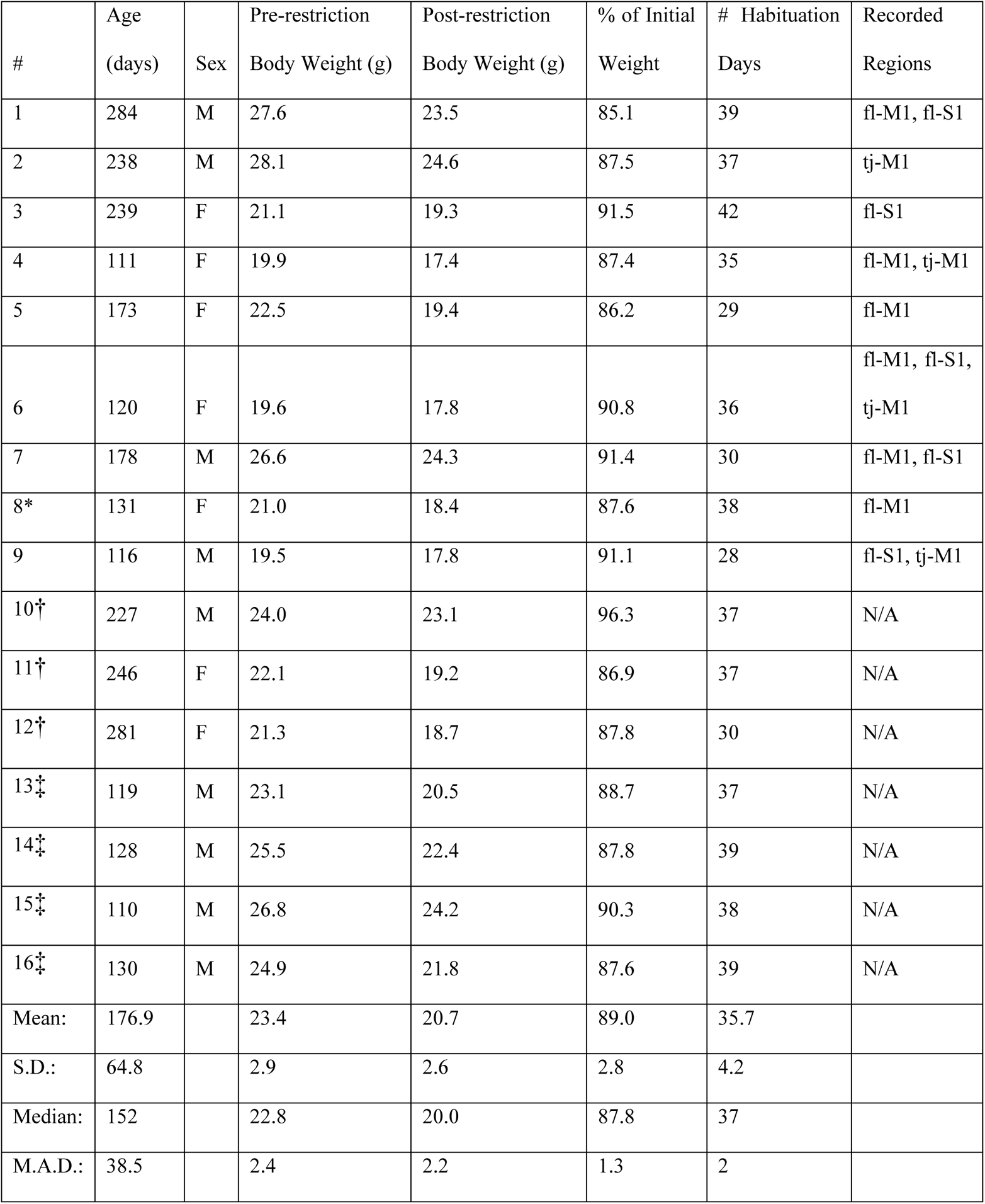
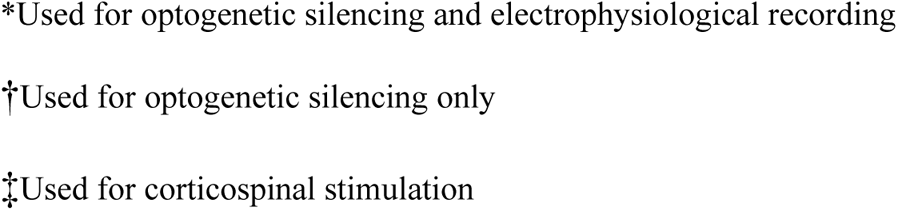
Mice appearing in this paper

| # | Age<br>(days) | Sex | Pre-restriction<br>Body Weight (g) | Post-restriction<br>Body Weight (g) | % of Initial<br>Weight | # Habituation<br>Days | Recorded<br>Regions |
| --- | --- | --- | --- | --- | --- | --- | --- |
| 1 | 284 | M | 27.6 | 23.5 | 85.1 | 39 | fl-M1, fl-S1 |
| 2 | 238 | M | 28.1 | 24.6 | 87.5 | 37 | tj-M1 |
| 3 | 239 | F | 21.1 | 19.3 | 91.5 | 42 | fl-S1 |
| 4 | 111 | F | 19.9 | 17.4 | 87.4 | 35 | fl-M1, tj-M1 |
| 5 | 173 | F | 22.5 | 19.4 | 86.2 | 29 | fl-M1 |
| 6 | 120 | F | 19.6 | 17.8 | 90.8 | 36 | fl-M1, fl-S1,<br>tj-M1 |
| 7 | 178 | M | 26.6 | 24.3 | 91.4 | 30 | fl-M1, fl-S1 |
| 8* | 131 | F | 21.0 | 18.4 | 87.6 | 38 | fl-M1 |
| 9 | 116 | M | 19.5 | 17.8 | 91.1 | 28 | fl-S1, tj-M1 |
| 10† | 227 | M | 24.0 | 23.1 | 96.3 | 37 | N/A |
| 11† | 246 | F | 22.1 | 19.2 | 86.9 | 37 | N/A |
| 12† | 281 | F | 21.3 | 18.7 | 87.8 | 30 | N/A |
| 13‡ | 119 | M | 23.1 | 20.5 | 88.7 | 37 | N/A |
| 14‡ | 128 | M | 25.5 | 22.4 | 87.8 | 39 | N/A |
| 15‡ | 110 | M | 26.8 | 24.2 | 90.3 | 38 | N/A |
| 16‡ | 130 | M | 24.9 | 21.8 | 87.6 | 39 | N/A |
| Mean: | 176.9 |  | 23.4 | 20.7 | 89.0 | 35.7 |  |
| S.D.: | 64.8 |  | 2.9 | 2.6 | 2.8 | 4.2 |  |
| Median: | 152 |  | 22.8 | 20.0 | 87.8 | 37 |  |
| M.A.D.: | 38.5 |  | 2.4 | 2.2 | 1.3 | 2 |  |

\*Used for optogenetic silencing and electrophysiological recording
†Used for optogenetic silencing only
‡Used for corticospinal stimulation

For the hand-hand distance, D, alignment to the peak of the regrip showed a roughly bell-shaped trajectory, with an amplitude of 3.2 ± 0.6 mm and duration (based on width at half-maximum of the peak) of 25.1 ± 4.6 ms (**Fig. 2B, middle**).

These basic features of oromanual events were highly stereotyped, as reflected by the low coefficients of variation for the kinematic parameters (**Table 2**). Oromanual event durations, measured from the end of the lowering-from-mouth movement to the beginning of the transport-to-mouth movement, averaged 2.48 ± 0.95 s. The number of regrips per oromanual event was 3.9 ± 1.5 (regrips/oromanual), and most events (91 ± 11% across all mice) had at least one regrip. The number of regrips increased with oromanual duration by an average of 1.2 ± 0.2 regrips per second spent in an oromanual event (linear mixed-effects regression of number of regrips on duration with mouse as grouping variable, *n* = 9 mice, *F*_174_ = 40.4, *p* = 1.8 × 10^-9^), and the latency to the first regrip averaged 170 ± 49 ms. The durations of each component were distributed in an approximately log-normal manner (**Fig. 2C**). Transitions between oromanual and holding occurred at a rate of 0.24 ± 0.06 Hz with 36 ± 8% of time spent in the oromanual mode.

**Table 2:**
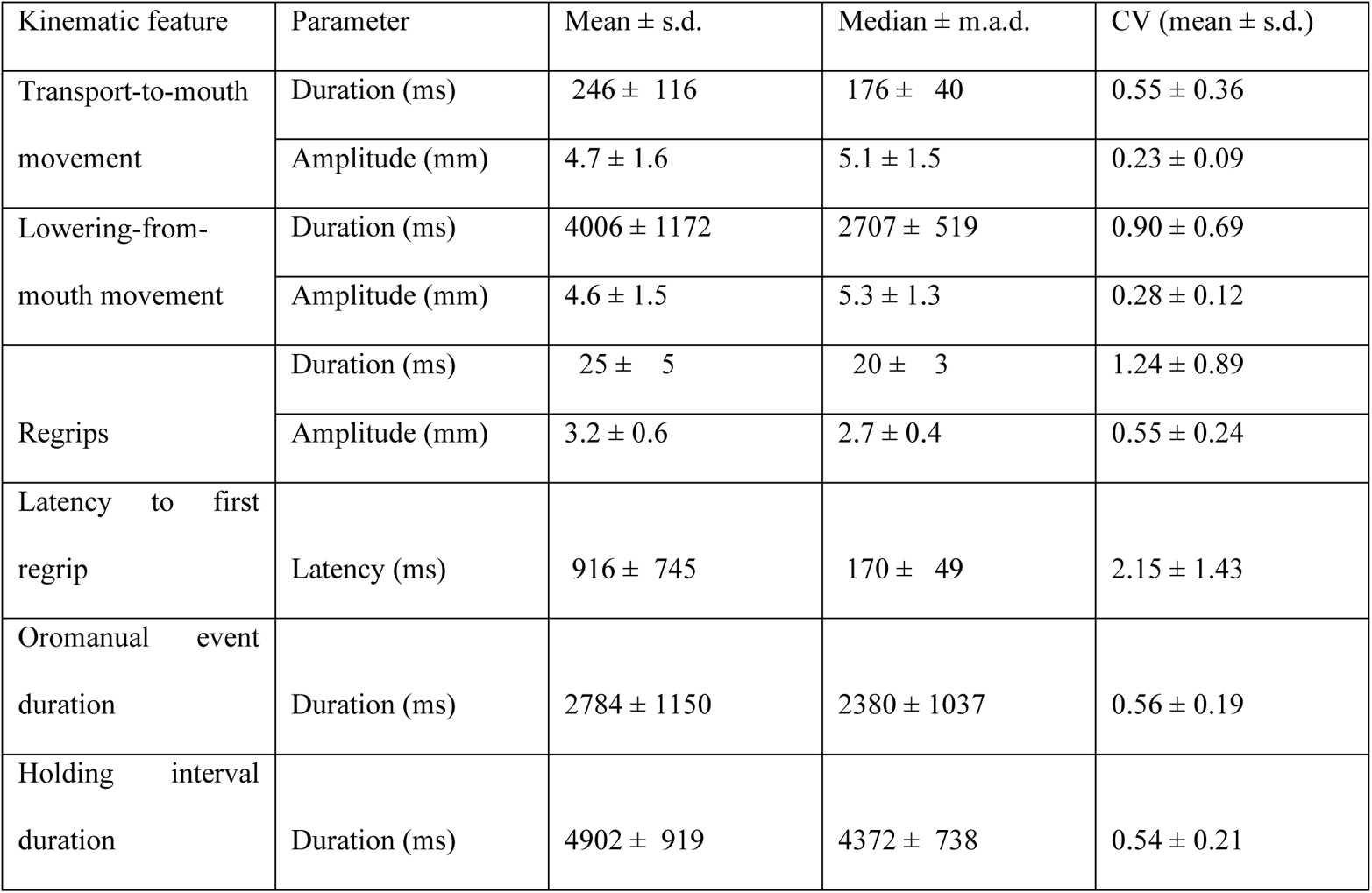
Kinematic properties of oromanual events

We used optogenetic silencing to assess the effects of transient motor cortex inactivation on food-handling in a cohort of Ai32xPV-cre mice (*n* = 4, **Table 1, Methods**) (Morandell and Huber, 2017; Li et al., 2019). During bilateral silencing of forelimb M1, regrip frequency fell compared to periods preceding and following silencing (repeated-measures ANOVA, *n* = 4, *F*_2_ = 6.7, *p* = 0.03) (**Fig. 2D, Video 3**). Other kinematic parameters were not significantly affected (trajectories of transport-to-mouth, lowering-from-mouth, or regrip movements; durations of oromanual or holding modes; likelihood of transitions).

Conversely, we used optogenetic stimulation of channelrhodopsin-2 (ChR2) expressing corticospinal neurons in forelimb M1 to assess how transient motor cortex activation might affect food-handling (*n* = 4 mice, **Table 1, Methods**). Selective corticospinal activation, when delivered during holding intervals, rapidly evoked transport-to-mouth movements (**Fig. 2E, Video 4**), with a significantly higher probability of observing a transport-to-mouth in the 400 ms following stimulus onset in holding (repeated-measures ANOVA, *n* = 4, *F*_2_ = 112.1, *p* = 1.8 × 10^-5^) compared to one second before (pre vs peri: *p* = 0.006, Bonferroni method) or after (peri vs post: *p* = 0.005). Regrip rate was not affected, nor were other kinematic parameters. This effect was not seen in one mouse where corticospinal transfection failed, resulting in no ChR2 expression, nor was it seen in PVxAi32 mice, ruling out the possibility that the evoked transport-to-mouth is a visual reaction to the laser.

Collectively, these analyses quantify the major kinematic features of oromanual events and establish that modulation of forelimb M1 activity influences food-handling behavior, providing a basis for detailed analysis of related cortical activity.

### Phasic-tonic oromanual-related activity in forelimb M1

To assess how the firing of active units related to the different features of oromanual events, we aligned the kinematic and neural data to the transport-to-mouth, regrip, or lowering-from-mouth movements. As shown in the example experiment (**Fig. 3A-F**), while the hand-nose distance, L, remained low throughout oromanual events (**Fig. 3A-B**), the example active unit increased in firing around transport-to-mouth movements (**Fig. 3C,E; left**) and regrips (**Fig. 3C,E; middle**), then fell to lower levels, returning to baseline with the lowering-from-mouth movement (**Fig. 3C,E; right**). This pattern was observed for many active units recorded in this experiment (**Fig. 3D-F**), and across mice (**Fig. 3G-I**).

**Fig. 3:**
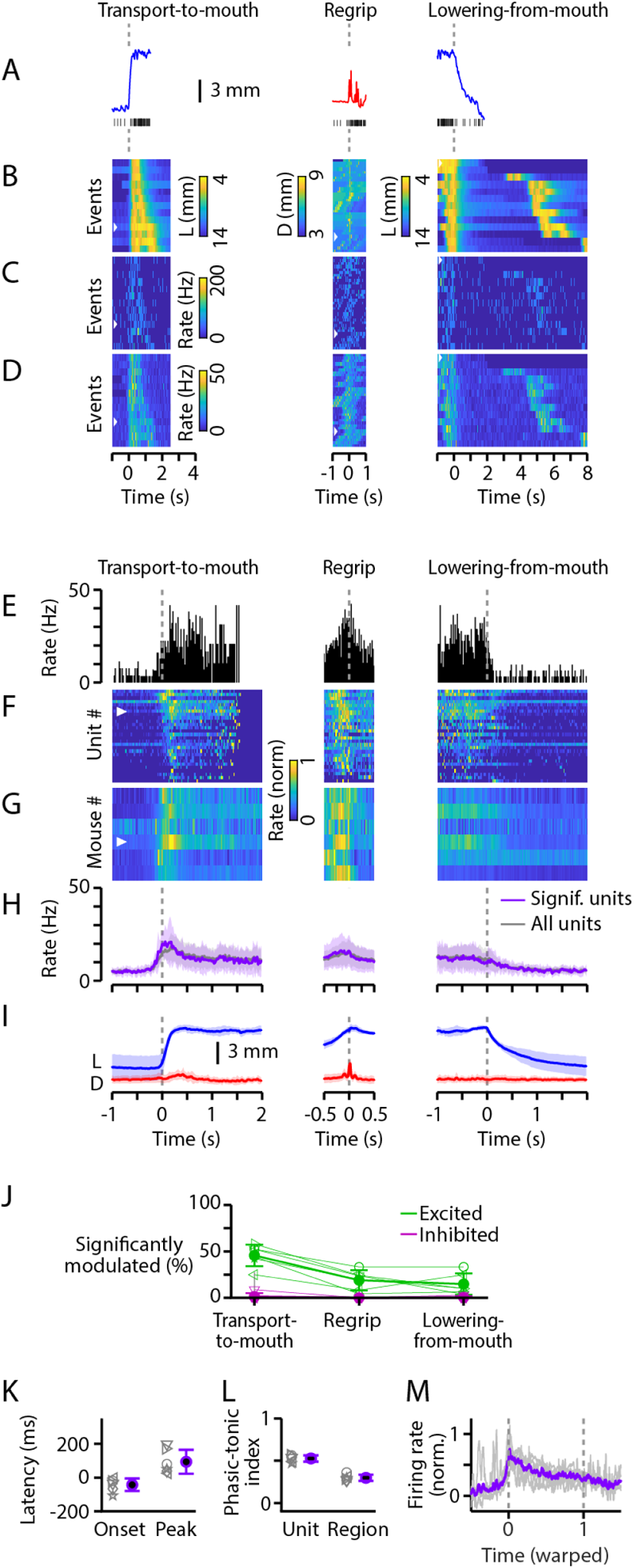
Phasic-tonic oromanual-related activity in forelimb M1. (A) Traces of L (blue, reverse y-axis) for an example transport-to-mouth movement (left), D (red) for an example regrip (middle), and L for an example lowering-from-mouth movement (right). (B) Heatmap of all peri-oromanual event traces of kinematics for the same experiment as in (A). Left: Map of L (inverse color scale), aligned to the transport-to-mouth onset, sorted by oromanual event duration. Middle: Map of D, aligned to regrips, sorted by latency. Right: Map of L, aligned to lowering-from-mouth onset and sorted by holding interval duration. White arrows denote the corresponding events in (A). (C) Same as (B) for an example forelimb M1 single unit. (D) Same as (B) for the average firing rate of all active units recorded from forelimb M1 in the example experiment. (E) Peri-event time histograms (PETHs) for the example single unit in (C). (F) Peak-normalized PETHs of all significantly modulated (see **Methods**) active units recorded in the same experiment, plotted as a heatmap. White arrow denotes the example single unit in (C) and (E). (G) Average peak-normalized event-aligned firing rates across significantly modulated forelimb M1 active units and across experiments for all mice with forelimb M1 recordings (*n* = 6). (H) Mouse-average event-aligned firing rates (shaded region, mean ± s.d.) for all units (grey) and only significantly modulated units (purple). (I) Mouse-average event-aligned L (blue, reverse y axis) and D (red) traces. (J) Percentages of forelimb M1 active units significantly excited (green) or inhibited (purple) for each event type. Thin lines are means over experiments for individual mice, thick lines and error bars are mean ± s.d. over mice. (K) Onset and peak latencies for active units in forelimb M1. Grey symbols are means over simultaneously recorded units and then experiments for individual mice, purple symbols and error bars are means ± s.d. over mice. (L) Phasic-tonic indices (PTIs, see **Methods**) for forelimb M1 active units (“Unit”) and average firing rate traces (“Region”). Grey symbols are means over experiments (after first averaging over simultaneously recorded units for Unit PTIs) for individual mice, purple symbols and error bars are mean ± s.d. over mice. (M) Normalized average forelimb M1 firing rate traces after time-warping each oromanual event to have the same duration. Thin lines are means over experiments for each mouse and thick lines are mean over mice.

Across experiments and mice, many active units in forelimb M1 were significantly positively modulated (bootstrap test, Holm-Bonferroni corrected *p*-values < 0.05, 36 ± 19% of active units, mean ± s.d., *n* = 6 mice, **Fig. 3J**) around the transport-to-mouth movement and none were significantly inhibited. Firing rate increases of significantly excited active units began 41 ± 33 ms before the transport-to-mouth movement and peaked 112 ± 96 ms after movement onset (**Fig. 3K**). Similarly, we analyzed how cortical activity relates to regrips (middle panels in **Fig. 3A-I**). A small fraction of active units was significantly excited around regrips (10 ± 12%), while very few (0.4 ± 1.2 %) were inhibited (**Fig. 3J**). Finally, very few active units were significantly modulated around the lowering-from-mouth movement (0.3 ± 1% excited, 1.9 ± 5.9 % inhibited; **Fig. 3J**). Because activity tended to fall to lower levels after an early peak, we calculated a phasic-tonic index (PTI) (Shalit et al., 2012) for all active units (not just those significantly modulated) as the ratio of (FR_peak_ – FR_end_)/(FR_peak_ + FR_end_), i.e. the difference between firing in a 200 ms window surrounding its peak to firing in a 200 ms window at the end of each oromanual event, divided by their sum. (Perfectly tonic activity thus yields a PTI of 0, whereas perfectly phasic activity yields a PTI of 1.) Across mice, the average PTI was 0.46 ± 0.08 (**Fig. 3L**), indicating a phasic-tonic pattern in which the average active unit’s firing peaks early and then decays to roughly one quarter of its peak by the end of the oromanual event.

Given the generally unidirectional modulation of active unit firing around oromanual events in forelimb M1, we next considered the probe-average firing rate, which showed a similar pattern for all units or only those significantly modulated (**Fig. 3G-H**). Across mice, activity started to rise slightly before the hands began to move towards the mouth, peaked around when the hands reached the mouth, and fell shortly thereafter, remaining elevated above baseline until the lowering-from-mouth movement. The average PTI for probe-average activity was 0.28 ± 0.05 (**Fig. 3L**), indicating a phasic-tonic pattern in which activity decays to about one half of its peak by the end of an oromanual event. This pattern was also apparent when oromanual events and the associated probe-average activity traces were time-warped to equal durations (**Fig. 3M**).

### Tonic activity in tongue/jaw M1 and an intermediate pattern in forelimb S1

Because oromanual movements also involve the mouth and jaw, we made additional recordings in an area designated “tongue/jaw M1”, implicated in oral movements and located anterior and lateral to forelimb M1 (Mayrhofer et al., 2019). Tongue/jaw M1 activity was overall higher during oromanual events compared to holding intervals (mean ± s.d.: holding 7.1 ± 4.9 Hz; oromanual 18.1 ± 10.0 Hz; paired *t*-test: *t*_3_ = 3.70, *p* = 0.03, *n* = 4 mice). As shown in the examples and borne out in averages (**Fig. 4A-I**), alignment to the onset of the transport-to-mouth showed a rise in activity as the hands raised towards the mouth, with 52 ± 25% of active units significantly excited and none significantly inhibited (**Fig. 4J**). Of the significantly excited units, their onset latency was 212 ± 223 ms after the transport-to-mouth movement onset and their peak latency was 484 ± 107 ms (**Fig. 4K**). A small fraction (16 ± 11%) of active units were significantly excited around regrips and 2.6 ± 2.9% significantly inhibited (**Fig. 4J**). Very few active units were significantly modulated around lowering-from-mouth movements (excited: 1.4 ± 3.2%, inhibited: 7.1 ± 8.4%; **Fig. 4J**). Tongue/jaw M1 units had a PTI of 0.23 ± 0.04 (**Fig. 4L**). Contrasting with forelimb M1, probe-average firing activity in tongue/jaw M1 activity remained elevated over the course of oromanual events, only decaying to baseline levels with the return to holding posture (**Fig. 4G-H**). This was reflected by a much lower phasic-tonic index of 0.002 ± 0.09 (**Fig. 4L**), which was not significantly different from zero (paired *t*-test: *t*_3_ = 0.32, *p* = 0.77), indicating tonic activity. A tonic pattern was also evident in the time-warped traces (**Fig. 4M**).

**Fig. 4:**
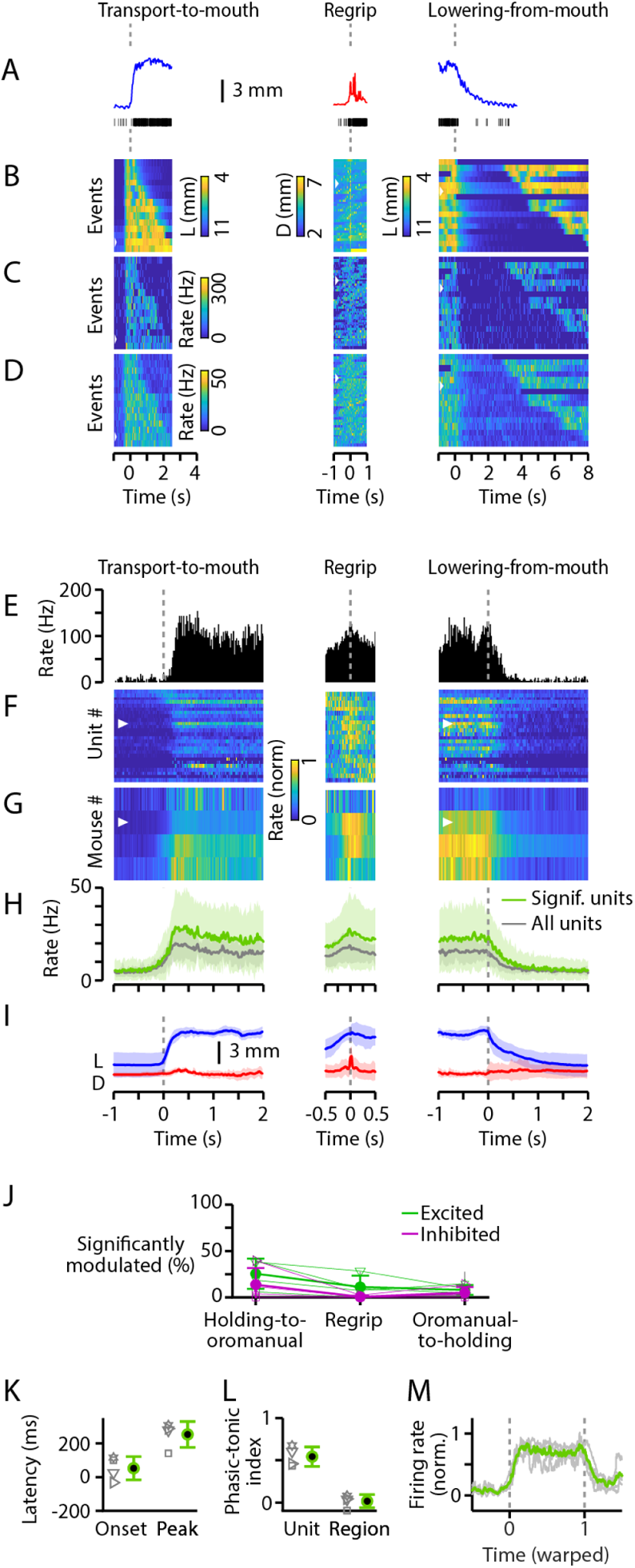
Tonic oromanual-associated activity in tongue/jaw M1. As Fig. 3, but for all tongue/jaw M1 recordings (*n* = 4).

Because oromanual movements also involve forelimb somatosensation, we also recorded from the forelimb region of the primary somatosensory (S1) area, located laterally adjacent to forelimb M1 (Yamawaki et al., 2021). Forelimb S1 activity was overall higher during oromanual events compared to holding intervals (mean ± s.d.: holding 7.9 ± 3.2 Hz; oromanual 18.5 ± 8.4 Hz; paired *t*-test: *t*_4_ = 2.9, *p* = 0.04, *n* = 5 mice). As shown in the examples and borne out in averages (**Fig. 5A-I**), alignment to the onset of the transport-to-mouth showed a rise in activity as the hands raised towards mouth, with 26 ± 19% of active units significantly excited but none significantly inhibited (**Fig. 5J**). Of the significantly excited units, their onset latency was 50 ± 106 ms after the transport-to-mouth movement onset and their peak latency was 286 ± 83 ms (**Fig. 5K**). There was a significant effect of cortical area on onset (Kruskal-Wallis ANOVA: *χ*^2^ = 6.7, *p* = 0.04) and peak latency (*χ*^2^ = 10.6, *p* = 0.005), with follow-up tests showing significantly longer onset (*p* = 0.03, Bonferroni method) and peak (*p* = 0.005) latencies in tongue/jaw M1 compared to forelimb M1.

**Fig. 5:**
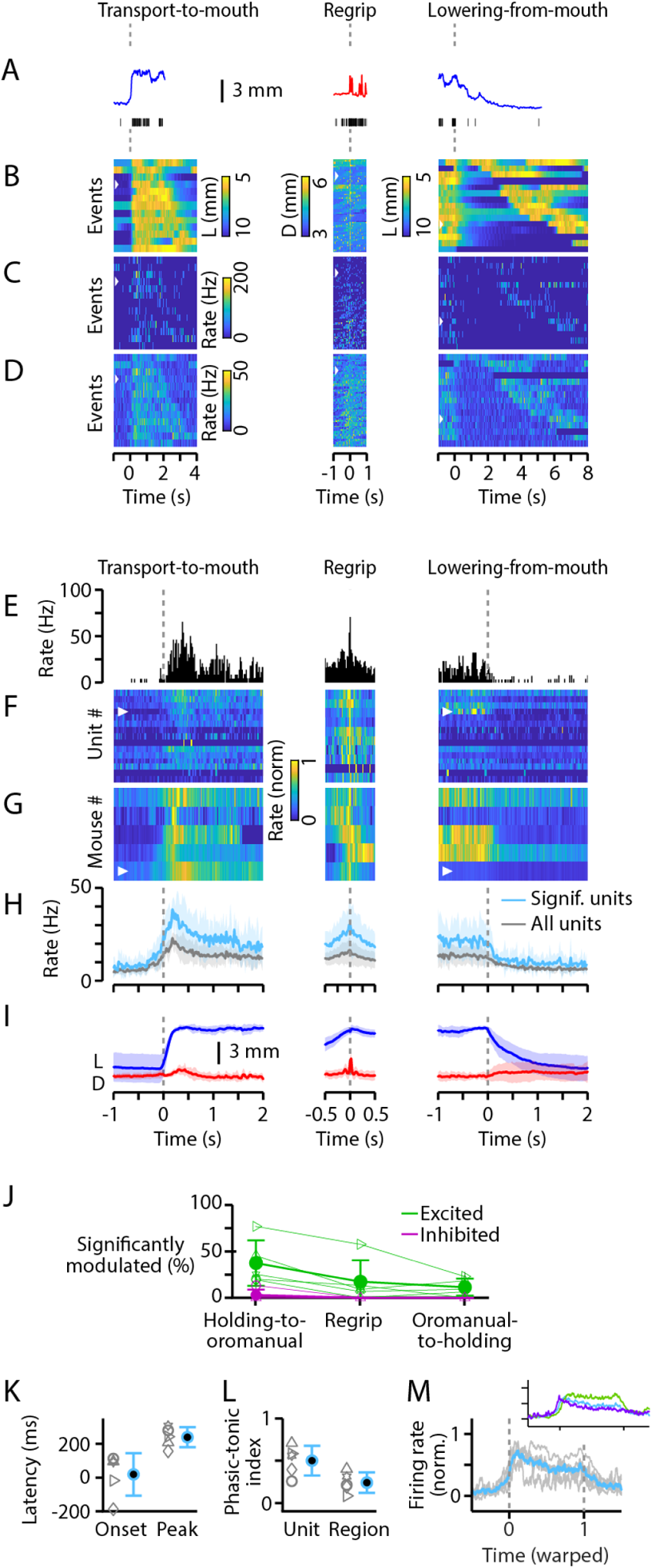
Tonic oromanual-associated activity in forelimb S1. As Fig. 3, but for all forelimb S1 recordings (*n* = 5). Inset in (N) shows mouse-average time-warped firing rate traces for all three regions on the same axis (purple, forelimb M1; green, tongue/jaw M1; teal, forelimb S1).

Aligning to regrips, 17 ± 12% of forelimb S1 active units were significantly excited but only 1.8 ± 4.9% significantly inhibited (**Fig. 5J**). Few active units were significantly modulated around lowering-from-mouth movements (11 ± 9%), and none were significantly inhibited (**Fig. 5J**). Forelimb S1 units had a lower PTI compared to those in forelimb M1 of 0.33 ± 0.17 (**Fig. 5L**). Probe-average forelimb S1 activity showed a similar phasic-tonic pattern to that seen in forelimb M1 (**Fig. 5G-H**), with a comparable phasic-tonic index of 0.22 ± 0.11 (**Fig. 5L**). There were significant effects of cortical area on phasic-tonic index regardless of whether they were calculated on active units or probe-average responses (permutation test, *p* < 2 × 10^-16^), with follow-up tests showing tongue/jaw M1 was significantly more tonic than forelimb M1 (*p* = 0.00002), but no difference between forelimb M1 and forelimb S1 (*p* = 0.08). Controlling for region, active units were significantly more phasic than probe-average activity (*p* = 0.0004). The interaction between region and PTI calculation method was not significant (*p* = 0.49). The temporal profile of both the time-warped activity for forelimb S1 (**Fig. 5M**) was intermediate between those of the other two regions.

These results indicate that cortical activity during food-handling exhibits both area-common and area-specific patterns. Activity was strongly associated with oromanual events in all three areas, reaching higher peak levels in tongue/jaw M1 and forelimb S1 than in forelimb M1. Oromanual-related activity followed a phasic-tonic pattern in forelimb M1, tonic pattern in tongue/jaw M1, and an intermediate pattern in forelimb S1. Timing of the initial rise was similar in all three areas, peaking earliest in forelimb M1 and slightly later in tongue/jaw M1 and forelimb S1.

### Phasic and tonic activity classes within and across areas

Averaging activity across probe channels may obscure heterogeneity in the patterns of event-aligned activity exhibited by active units within a given area. To address this, we used non-negative matrix factorization (NNMF) (Lee and Seung, 1999) to simultaneously perform dimensionality reduction and unsupervised clustering (**Methods**). Unlike related dimensionality reduction techniques such as principal components analysis, NNMF constrains the extracted features to be non-negative, making it particularly suited to spike train data, and also clusters the data by assigning each neuron to a cluster based on the factor for which that neuron has the greatest weight (Xu et al., 2020).

Applying NNMF to pooled data from all three areas, where the number of clusters was chosen automatically by bi-cross-validation, revealed two clusters of activity (**Fig. 6**). One cluster followed a phasic-like pattern, rising before the transport-to-mouth movement, briefly peaking, and falling thereafter. A second cluster, by contrast, showed a tonic-like pattern delayed relative to the transport-to-mouth movement, peaking around regrips, and remaining elevated until the lowering-from-mouth movement. These phasic-like and tonic-like patterns pertain to the clusters as a whole and not to the active units themselves, whose activity exhibited various patterns and peaked at various times in relation to the kinematics (**Fig. 6A-D**), and were more phasic than probe-average activity (**Fig. 3M, Fig. 4M, Fig. 5M**). Further, these clusters were differentially apportioned across cortical areas (**Fig. 6E**), creating the distinct area-specific average activity patterns. Applying NNMF to data from each area individually also found two clusters in each area (**Fig. S2**), which were qualitatively similar to the clusters found in the pooled data, although with a stronger transport-to-mouth peak in the forelimb M1 phasic-like cluster and a strong regrip-aligned peak in the forelimb S1 tonic-like cluster.

**Fig. 6:**
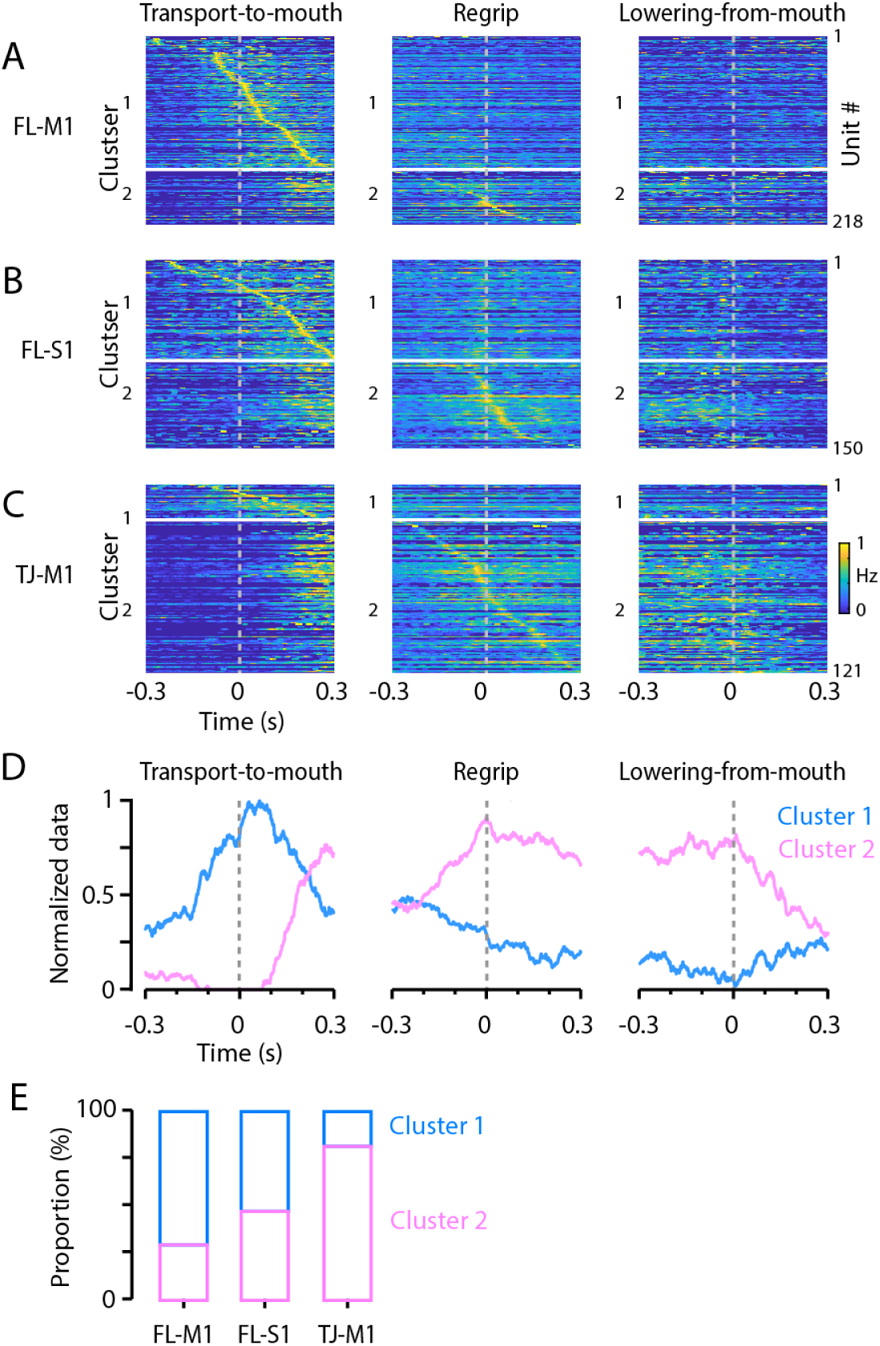
Phasic and tonic activity classes within and across areas. (A) Forelimb M1 unit activity normalized to maximum firing rate, aligned to transport-to-mouth (left), regrip (middle), and lowering-from-mouth (right) movements, and sorted by time of peak firing around transport-to-mouth movements for cluster 1 and regrips for cluster 2. Neurons were assigned to two clusters by applying NNMF to pooled data from all three areas. (B, C) Same as (A), for forelimb S1 and tongue/jaw M1, respectively. (D) NNMF factor weights for the two clusters. (E) Proportions of cluster 1 and 2 units for each area.

### Forelimb areas predict future hand position while tongue/jaw M1 encodes current hand position

The preceding analyses maybe biased by the choice of kinematic features considered and events aligned to. As a complementary approach, we correlated cortical activity with the 3D positions of the hands using whole (un-event-aligned) recordings. To do so we fit general linear models (GLMs) from sliding windows of binned neural activity to the kinematics (**Fig. 7**), using ridge regularization and cross-validation to deal with overfitting and multicollinearity (see **Methods**). Initially, the decoding window was large, using both past and future spiking activity to predict kinematics.

**Fig. 7:**
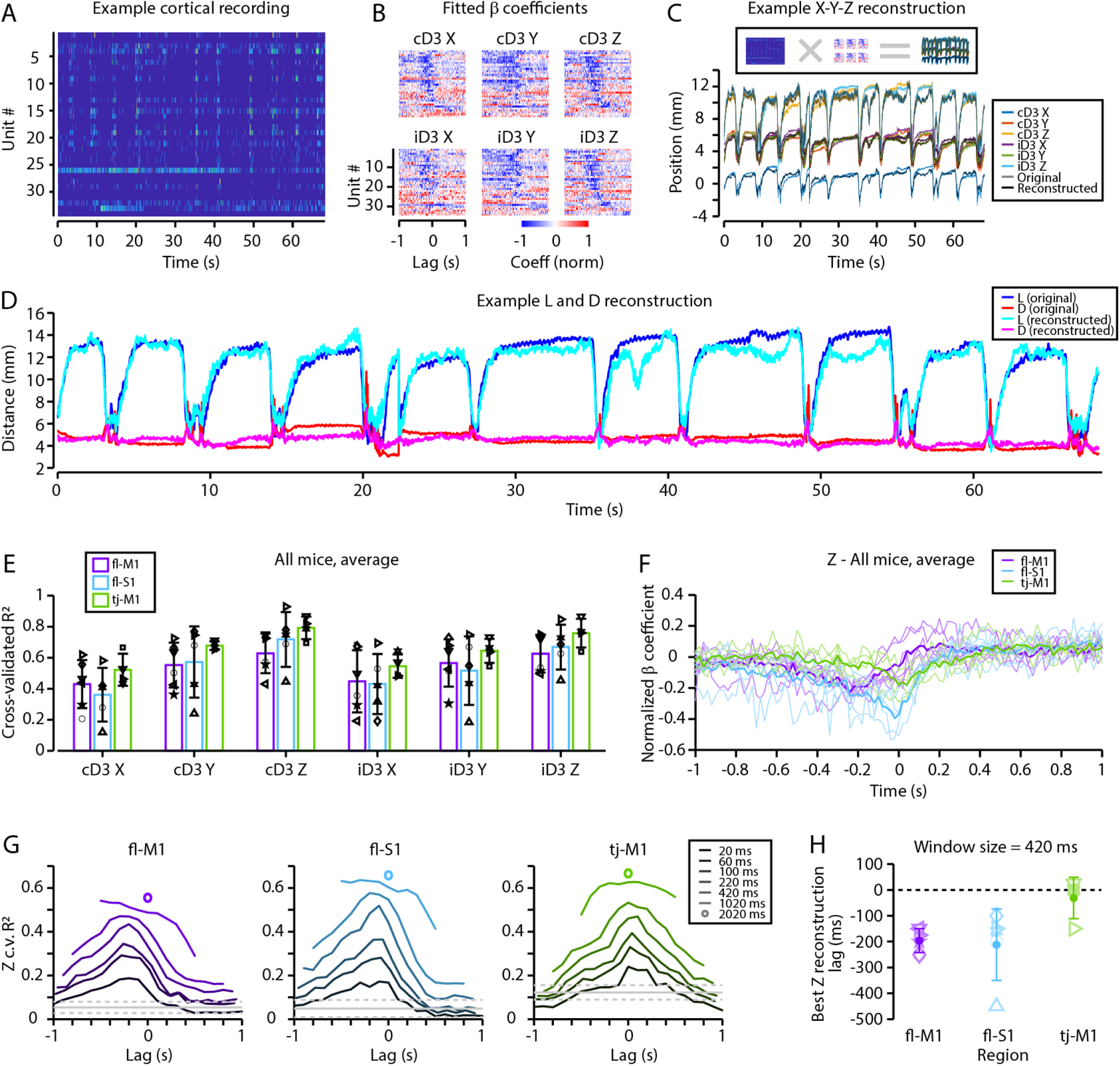
Hand position during food handling can be accurately decoded from cortical spiking activity. (A) Example raster showing all active units from a forelimb M1 recording. (B) To reconstruct kinematics, a ridge-regularized general linear model (GLM) was fit to the X, Y, and Z coordinates of each hand (cD3: contralateral third digit (D3), iD3: ipsilateral D3) using 2.02-second sliding windows of firing activity. The corresponding fitted GLM coefficients from the recording in (A), normalized to range [−1, 1] are shown. Active units are ordered by time-to-trough of the contralateral D3 Z coefficients. (C) X, Y, and Z hand trajectories (lighter colors) for the same example recording shown in (A-B) and the corresponding reconstructed trajectories (darker colors). (D) L (blue) and D (red) traces from the recording in (A-C) compared to L (cyan) and D (magenta) traces calculated from the reconstructed hand trajectories. (E) Cross-validated reconstruction accuracy (c.v. *R*^2^) for each coordinate and region (purple: forelimb M1, teal: forelimb S1, green: tongue/jaw M1). Error bars are mean ± s.d over mice. Symbols are averages over experiments for individual mice. (F) GLM coefficients normalized to the range [−1,1] for the Z-coordinate GLMs averaged over neurons, experiments, hands, and mice for forelimb M1 (purple), forelimb S1 (teal), and tongue/jaw M1 (green). Thin lines are individual mice, thick lines are mean over mice. (G) Mouse-average c.v. *R*^2^ for the Z-coordinate (averaged over ipsi- and contralateral hands) when varying the size and central lag of the window used for reconstruction. Solid grey lines indicate reconstruction accuracy expected by chance, dashed lines are mean ± 2 s.d. over shuffles of the chance reconstruction accuracy. Data for all coordinates are shown in Fig. S3. (H) Lag giving the highest hand-average Z-coordinate c.v. *R*^2^ for the 420 ms window as a function of region. Symbols are averages over experiments for individual mice, error bars are mean ± s.d. over mice.

As shown in an example (**Fig. 7A-B**) of a neural recording and the corresponding GLM coefficients for each hand coordinate, most active units showed a strong anticorrelation with kinematics at negative lags. The GLMs accurately reconstructed the 3D position of both hands in this recording (**Fig. 7C**), and hence also accurately reconstructed L (*R*^2^ = 88%) and D (*R*^2^ = 45%, **Fig. 7D**), even though the GLMs were not explicitly trained to do so. Across experiments, mice, and regions, reconstruction accuracy was high (**Fig. 7E**), with a significant effect of dimension (X, Y, or Z, perpendicular to the sagittal, coronal, and horizontal planes, respectively) on reconstruction accuracy (mixed ANOVA, side and dimension as within-subjects factors, region as between-subjects factor, *F*_2/24_ = 46.9, *p* = 1.0 × 10^-8^, Greenhouse-Geisser corrected), but not side (contra vs ipsi), region, or any of their interactions. Reconstruction accuracy was highest for Z and lowest for X (*p* < 0.05, Bonferroni corrected, for all pairwise follow-up comparisons).

Plotting the neuron-average GLM coefficients (**Fig. 7F**) shows the temporal relationship between activity and reconstructed kinematics. For forelimb M1, neurons are on average anticorrelated with forearm Z position at negative lags, peaking at around 200 ms, then very slightly positively correlated at short positive lags. Forelimb S1 is increasingly anticorrelated at negative lags approaching zero, whereas the strongest anticorrelation for tongue/jaw M1 is around zero lag. Similar results were seen for X and Y. To further explore the temporal relationship between activity and kinematics, we varied the size and central lag of the window used for fitting the GLMs (**Fig. 7G, Fig. S3**). Reconstruction accuracy increased monotonically with window size but even using a single bin was above chance at many lags. Reconstruction accuracy varied depending on the central lag of the reconstruction. To summarize this relationship, we considered the highest reconstruction accuracy lag for the 420 ms window (as for longer windows the temporal relationship was less clear). For both forelimb M1 and S1, accuracy peaked around 200 ms preceding kinematics, although forelimb S1 was more variable, whereas tongue/jaw M1 peaked around zero lag (**Fig. 7H**). There was a significant effect of region on best reconstruction lag (Kruskal-Wallis ANOVA, *χ*^2^ = 6.8, *p* = 0.03), with the only significant difference in follow-up tests being between forelimb M1 and tongue/jaw M1 (*p* = 0.03, Bonferroni method).

Together, these results confirm that significant information about forelimb position is carried in cortical firing activity during food handling. In forelimb M1 and S1, this information is predictive, leading kinematics, whereas in tongue/jaw M1 there is a tight temporal correlation between firing activity and hand position. This corroborates the earlier findings, in particular the better reconstruction accuracy for Z accords with the significant modulation of all areas by transport-to-mouth movements, and the zero-lag peak for tongue/jaw M1 fits with its tonic activity profile.

## DISCUSSION

We studied cortical activity associated with food-handling, as a step towards understanding the neurobiology of this ethologically critical behavior. The main findings show that cortical activity is (i) associated with specific kinematic features of food-handling movements relating to oromanual manipulation events, (ii) is distributed across multiple motor and somatosensory areas, with both area-common and area-specific features, (iii) exhibits prominent overall population-wide manipulation-related fluctuations but with some degree of heterogeneity at the level of individual neurons, and (iv) is temporally differentiated and predictive of current or future hand kinematics, in an area-dependent manner. Collectively our results provide a detailed characterization of multi-areal cortical activity associated with this form of manual dexterity in a food-handling mammal.

Our technical approach entailed addressing several methodological considerations relevant to food-handling behavior. Food-handling involves high-speed movements with millisecond-scale features, necessitating a video camera with high frame rate, sensitivity, and storage capacity. Dual camera views helped to mitigate problems of intermittent occlusion of visibility and provided datasets for 3D reconstruction of movement trajectories. Multichannel electrophysiology enabled spiking activity of cortical neurons to be sampled on a fast time scale comparable to the kinematic data. Regarding the behavioral analyses, most of these focused on experimenter-chosen features (holding vs oromanual modes, transport-to-mouth, lowering-from mouth, regrips), which potentially introduces bias. The GLM analysis avoided this by focusing on whole kinematic and neural traces, but newer unsupervised methods (Wiltschko et al., 2015; Batty et al., 2019; Pereira et al., 2020; Hsu and Yttri, 2021; Whiteway et al., 2021) might reveal further features or structure of food-handling behavior overlooked here.

The widespread cortical activity during the active, oromanual periods of food-handling is striking both for its spatial distribution and the depth of modulation, with large fractions of neurons in each area displaying significant manipulation-specific activity. Manipulation-related neural activity could (in principle, and in the extreme) be either heterogenous, with no overall pattern, or homogenous, with neurons covarying together as an ensemble. The active unit, probe-average, and reconstruction analyses suggest the second possibility. In aggregate, cortical activity varied unidirectionally with food-handling, increasing during active manipulation events and decreasing during passive holding intervals. However, two results show that this is an oversimplification. Active units in all regions are on average more phasic than the corresponding average traces, which would be impossible if they fired together as one homogenous population. Second, NNMF clustering revealed two activity patterns shared between areas, rather than one global pattern or a single cluster per area, implying deeper structure to the population response that is obscured by simple averaging.

Our results reveal both area-common and area-specific features of oromanual-associated cortical activity. Forelimb M1 showed a phasic-tonic activity pattern, peaking around the transport-to-mouth movement then decaying to above baseline firing rates. Forelimb M1 units were predictive of future movements and increased their firing tens of milliseconds before movement initiation, longer than reported latencies to electromyographic responses from mouse motor cortical stimulation (Ayling et al., 2009). Our and others’ silencing results (Guo et al., 2015; Mohan et al., 2021) argue against forelimb M1 directly driving food-handling movements, but suggest forelimb M1 does influence food-handling behavior. We interpret the lack of complete abolishment of food-handling movements by M1 silencing to reflect that other circuits in a distributed network are involved in supporting this behavior. Indeed, redundancy in the network controlling such a fundamental behavior would be highly adaptive, and has been seen in other motor behaviors (Li et al., 2016; Morandell and Huber, 2017). Stimulation of forelimb M1 corticospinal neurons evoked transport-to-mouth movements, but at sufficiently long latencies to suggest that the effect occurs via these downstream circuits rather than direct drive of spinal circuits alone. The precise circuits responsible remain an open question, but could involve corticospinal branches to the striatum, midbrain, or brainstem (BICCN, 2021; Nelson et al., 2021). In additional to forelimb M1, we also found manipulation-related activity in two other cortical regions, tongue/jaw M1 and forelimb S1. Activity in tongue/jaw M1 was delayed and tonic compared to forelimb M1. Rather than predicting future kinematics, it closely tracked the current hand position. This may be an epiphenomenon, with tongue/jaw M1 activity relating to orofacial movements during oromanual events that we were unable track due to occlusion by the hands (but may be revealed by electromyography of orofacial muscles such as the masseter). Alternatively, tracking the current forelimb position may facilitate the intricate coordination of the hands and mouth involved in food-handling. Similarly, some component of activity in forelimb M1 may represent information about current orofacial movements. Together with the lack of tongue/jaw M1 activity during the holding interval despite robust jaw activity as the mouse chews, this accords with the concept of an ethological action map in motor areas (Graziano, 2016), rather than simple somatotopy, as suggested by some recent motor mapping studies (Hira et al., 2015; Mercer Lindsay et al., 2019). Activity in forelimb S1 was quantitatively intermediate between forelimb M1 and tongue/jaw M1. This may be due to feed-forward input from forelimb M1 followed by incoming tactile information (Umeda et al., 2019) resulting in a temporally smeared version of the forelimb M1 activity trace. Alternatively, it may imply that forelimb S1 relays information from forelimb M1 to tongue/jaw M1. Characterization of connectivity between these regions and area-specific optogenetic stimulation will help distinguish these possibilities.

Our findings substantially advance knowledge about cortical involvement in manual dexterity. More detailed activity-perturbation studies and cell-type resolved methods have potential to elucidate the precise roles of cortical and subcortical circuits in this behavior. The population-wide fluctuations observed suggest neuromodulatory systems, particularly noradrenergic input from the locus coeruleus, as an avenue for future study. Finally, independent feeding is an important activity of daily living that may be lost as a result of conditions such as stroke and spinal cord injury. Deeper understanding of the neurobiology of food-handling, including similarities and differences across species, will inform efforts to restore function to such patients, and the results and methodological advances presented here contribute towards this goal.

## SUPPLEMENTARY FIGURES

**Fig. S1:**
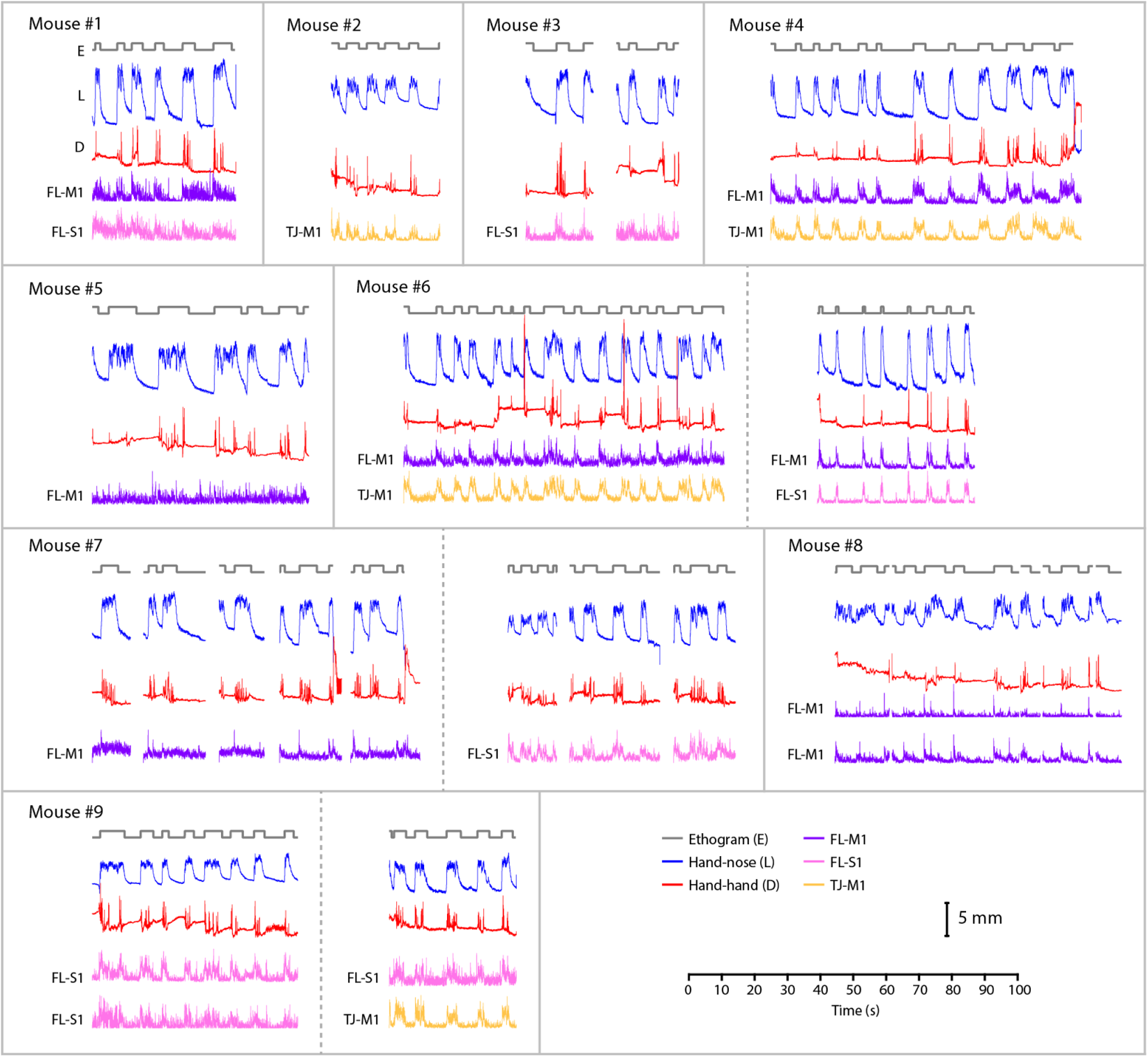
Traces of kinematic parameters and firing rates. Traces of kinematic parameters and firing rates are shown for one or more representative recording sessions for each mouse. Each recording session is of the handling of one food item. Mice were often recorded over multiple days (indicated by vertical dashed lines), each with multiple sessions. Probes were reinserted anew on subsequent days.

**Fig. S2:**
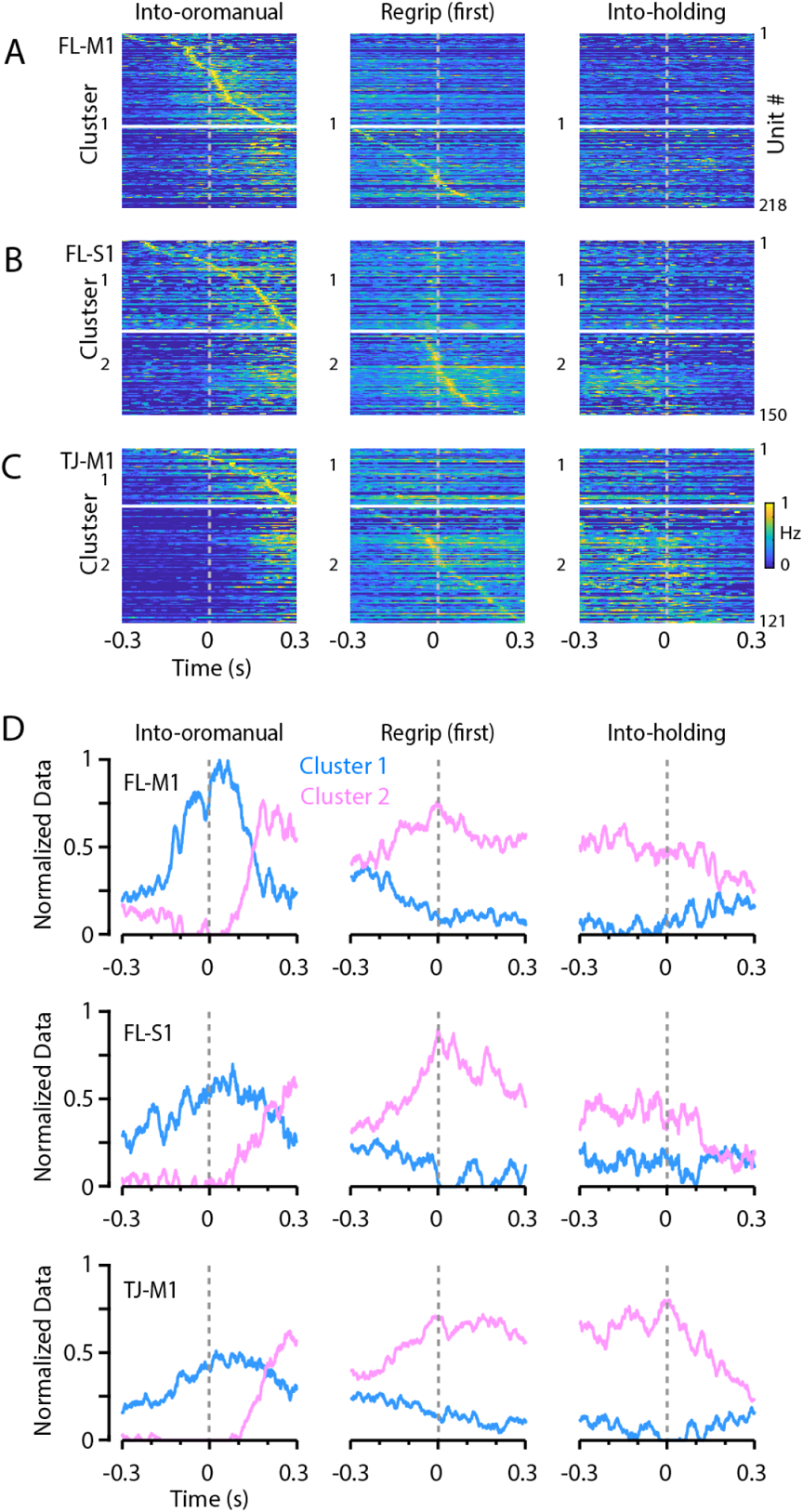
Phasic and tonic activity classes within and across areas, based on applying NNMF to data from each area individually. (A) Forelimb M1 unit activity, aligned to the transport-to-mouth movement (left), first regrips (middle), and lowering-from-mouth movement (right). Data were normalized to each unit’s maximum average firing rate and sorted by time of peak firing around the transport-to-mouth movement for cluster 1 and time of peak firing around the first regrip for cluster 2. Two clusters identified by NNMF are indicated. (B) Same as (A), for forelimb S1. Units were sorted by time of peak firing around the transport-to-mouth movement (C) Same as (A), for tongue/jaw M1. (D) Plots showing the NNMF factor weights for the clusters found in forelimb M1 (top), forelimb S1 (middle), and tongue/jaw M1 (bottom).

**Fig. S3:**
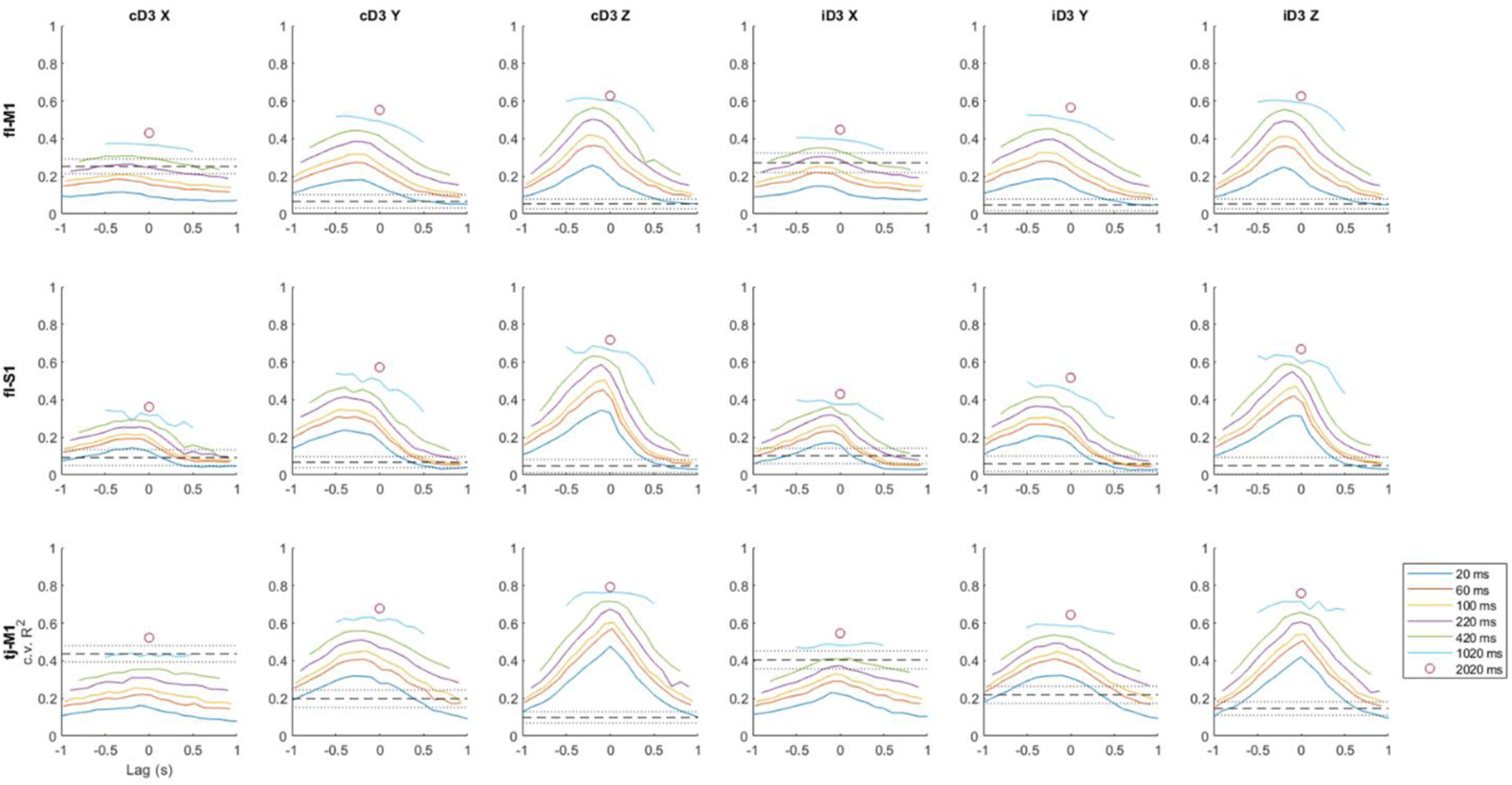
Additional GLM modeling results. Plots show the c.v. *R*^2^, averaged over mice, for the X-, Y-, and Z-coordinates of the ipsilateral (i-) and contralateral (c−) third digits (D3), for each of the three cortical areas, when varying the size and central lag of the window used for reconstruction. Dashed lines indicate reconstruction accuracy expected by chance, dotted lines are mean ± 2 s.d. over shuffles of the chance reconstruction accuracy.

## VIDEOS

**Video 1:** Example video segment featuring two oromanual events.

**Video 2:** Part of Video 1 slowed down to one-tenth speed, with tracking added. The DeepLabCut-tracked position of the nose is indicated by blue circles and each hand by red circles. The derived hand-hand (D) and hand-nose (L) distances are indicated by red and blue lines, respectively.

**Video 3:** Example oromanual event with optogenetic silencing, slowed to one-tenth speed. Current behavioral mode, laser state, and regrips are noted.

**Video 4:** Example spontaneous transport-to-mouth movement, slowed to one-tenth speed, followed by a transport-to-mouth movement evoked by corticospinal stimulation. Movement and laser onsets are noted.

## MATERIALS AND METHODS

### Animals

This study used experimentally naïve mice on a C57BL/6 background (stock no. 000664, The Jackson Laboratory, Bar Harbor, Maine) aged 111-284 days postnatal and weighing 17.4-24.6 g at the time of recording (19.5-28.1 g before food restriction, **Table 1**). For optogenetic silencing experiments, Ai32xPV-cre mice were generated by crossing homozygous B6.Cg-Gt(ROSA)26Sortm32(CAG-COP4*H134R/EYFP)Hze/J cre-dependent channelrhodospin2 (ChR2) reporter mice (stock no. 024109, The Jackson Laboratory) (Madisen et al., 2012) with homozygous B6.129P2-Pvalbtm1(cre)Arbr/J mice (stock no. 017320, The Jackson Laboratory) (Hippenmeyer et al., 2005) that express cre in parvalbumin (PV) expressing cells. Ai32xPV-cre mice thus express ChR2 in all PV-positive cells, including PV-positive interneurons in cortex, and so can be used for local silencing of cortex (Li et al., 2019). Mice of both sexes were used, consistent with NIH policy on sex as a biological variable in basic research. Mice were bred in-house, housed in groups with a 12 hour reverse light/dark cycle, and had free access to food and water prior to food restriction (see below). All experiments were conducted during the dark phase of the mice’s light cycle. Mice were used as they became available. As many brain regions as possible (of the total of six bilateral representations of the three areas of interest) were recorded from each mouse, hence no randomization to cohorts was necessary. All studies of mice were approved by the Northwestern University IACUC and fully complied with the animal welfare guidelines of the National Institutes of Health and Society for Neuroscience.

### Surgical procedures

#### Head-bar mounting

Under deep isoflurane anesthesia, mice were placed in a stereotaxic frame (Model 900, David Kopf Instruments, Tujunga, CA) and a ~1 cm^2^ circular incision was made to expose the cranium. The periosteum was removed and a titanium head-fixation bar (0.875 × 0.187 inches, cut by water-jet from 0.08 inch Ti-6Al-4V sheet, Big Blue Saw, Atlanta, GA) was placed on top of lambda, perpendicular to the central suture, and affixed using dental cement (C&B Metabond, Parkell, Edgewood, NY). The incision was then sutured to close the wound margins and cover any exposed cranium not covered by dental cement and/or the head-bar. Mice were given 0.3 mg/kg buprenorphine preoperatively and 1.5 mg/kg meloxicam postoperatively as analgesia, followed by a second dose of meloxicam 24 hours after surgery. Mice were single-housed following head-bar mounting.

#### Retrograde labeling of corticospinal neurons

For optogenetic stimulation of corticospinal neurons, pAAVretro-syn-ChR2(H134R)-GFP (#58880, Addgene, Watertown, MA) (Tervo et al., 2016) was injected into the spinal cord as previously described (Yamawaki et al., 2021) at the same time as head-bar mounting. Laminectomies were performed at cervical level 6. Injection pipettes were fabricated from glass capillary micropipettes (Wiretrol II, Drummond Scientific Company, Broomall, PA) using a pipette puller (PP-830, Narishige, Tokyo, Japan) and beveled to a sharp edge with a microgrinder (EG-400, Narishige). Pipettes were back-filled with mineral oil, tip-filled with virus, and advanced to the spinal cord using a 3-axis digital manipulator (51906, Stoelting, Wood Dale, IL). The dura was punctured and virus injected at 10 nL/min 0.4 mL lateral to midline at depths of 0.6 and 0.4 mm using a one-axis oil hydraulic micromanipulator (MO-10, Narishige) to a total volume of 80 nL.

#### Craniotomies for linear arrays

One day prior to recording, mice were deeply anaesthetized with a cocktail of 80–100 mg/kg ketamine and 5–15 mg/kg xylazine injected intraperitoneally. A craniotomy or craniotomies were opened over the area(s) to be recorded using a dental drill (EXL-M40, Osada, Los Angeles, CA). The cortical surface was covered with Kwik-Sil (World Precision Instruments, Sarasota, FL) and mice were allowed to recover.

### Behavioral training

At least 3 days post-surgery, mice were food restricted to motivate feeding behavior. Mice were fed a measured amount of standard rodent diet each day to maintain their weight between 85 and 90% of pre-restriction body weight. At the time of recording mouse weights were 85.1-91.5% of initial body weight. Mice were monitored throughout the food restriction period for signs of ill-health (Guo et al., 2014), and body condition scores (Ullman-Cullere and Foltz, 1999) were taken each day. No signs of ill-health were observed and no mouse fell below a body condition score of 3 throughout the study.

The experimental apparatus comprised a raised platform on which a 3D-printed hut was placed for the mouse to sit in, head-bar holders with screw clamps, and a pellet dispensing tube. The hut was designed to have an arched profile to facilitate the hunched posture typically adopted by mice while eating by providing more room for the back to arch, while also being easy to 3D-print. It also incorporated an arm bar for the mice to rest their hands on when not eating. The height of the platform was adjustable to enable a comfortable posture for each mouse. Accordingly, the position of the arm bar below the head varied from mouse to mouse and in some cases during holding intervals (particularly those of long duration) mice would rest the forearms on the arm bar. Mice generally did not use the arm bar for support during bimanual oromanual events, however.

Starting 1 to 3 days after beginning food restriction, mice were familiarized with the experimenter and head-fixation apparatus following standard procedures (Guo et al., 2014). Briefly, mice were first acclimatized to handling by the experimenter, then introduced to the experimental apparatus. Food rewards (20 mg dustless precision grain pellets, Bio-Serv, Flemington, NJ) were presented to the mice from the dispensing tube. Once mice consistently ate from the tube, they were introduced first to gentle head-fixation by hand, then to full head-fixation. Video and electrophysiological recordings were taken after four to six weeks of habituation, by which time mice were able to comfortably and consistently retrieve, handle, and consume pellets from the dispenser while head-fixed. For experimental recordings, the head-fixed mice were given black oil sunflower seeds (shells removed) or large grain pellets (45 mg, Bio-Serv), presented by spoon, as these larger food items facilitated longer duration recordings.

### Videography and kinematic analysis

Videos were obtained with a high-speed CMOS-based monochrome video camera (Phantom VEO 710L, Vision Research, Wayne, NJ). Videos were acquired at 1000 frames per second (fps), 999.6 µs exposure time, and 1024 × 512 pixel field of view. Two oblique views of the mouse were obtained by mounting two 50 × 50 mm flat enhanced aluminum surface mirrors (#43-876, Edmund Optics, Barrington, NJ) and a 50 mm anti-reflection coated equilateral prism (#49-435, Edmund Optics) in the camera optical path. A prime lens (Nikon AF Micro-NIKKOR 60mm f/2.8D, Nikon, Tokyo, Japan) was mounted on the camera body. The mouse was illuminated from both sides and slightly below using two red LEDs (M660L1 and MLEDC25, ThorLabs, Newton, NJ). Camera and video recording settings were controlled with Phantom Camera Control Application v3.5 (Vision Research). Video was recorded to the camera memory and then saved to disk as uncompressed Phantom Cine files, later converted to H.264-encoded MP4 files. Video recording was triggered by a TTL pulse delivered from an NI USB-6229 data acquisition board (National Instruments, Austin, TX).

Videos were cropped to isolate each view using ffmpeg (ffmpeg.org) and then markerless tracking of the nose, digits, and jaw was performed using DeepLabCut (Mathis et al., 2018) as described (Barrett et al., 2020). From these two sets of 2D trajectories for each body part, 3D trajectories were reconstructed using Anipose (Karashchuk et al., 2020). Anipose’s camera model was calibrated for each experimental session using videos of a ChAruCo board at various angles captured at the end of the session without adjusting the lens settings.

### Ethogramming

Using the DeepLabCut-extracted and Anipose-reconstructed trajectories, each video was temporally parsed into holding, oromanual, and other postural modes based on the distance between the third digit (D3) of each hand and the nose. Two thresholds were set for each hand, one dividing the oromanual events from holding intervals, and one dividing holding intervals from non-food-handling related behavior (e.g. resting the hand on the arm bar). Video frames were assigned to holding intervals when both hands were in holding and to oromanual events when both were in oromanual. In the case of a slight delay in one hand crossing the oromanual threshold relative to the other, the threshold crossing time was set to the mean of the threshold crossing times for the two hands. Brief mode transitions due to fluctuations about the threshold were removed. Thresholds for each hand and minimum mode duration for inclusion were set manually per video based on visual inspection of the hand-nose distance traces and resulting ethogram. Periods of unimanual behavior (one hand in holding/oromanual and the other at rest) were uncommon and excluded from analysis, as were periods where both hands were at rest or where the tracking was poor.

Having constructed a rough three-mode ethogram (oromanual/holding/other) based on threshold crossings, we then calculated the hand-hand distance D as the distance between the D3s of each hand and the bimanual hand-nose distance L as the distance between the nose and the midpoint of both D3s. The latter was used to more precisely determine the timing of each transition movement (transport-to-mouth and lowering-from-mouth) by fitting model functions to the corresponding L trace. For each threshold crossing, the segment of the L trace from halfway to the previous threshold crossing to halfway to the next crossing was taken for fitting. Model functions were chosen to approximate the characteristic shape of each movement. Transport-to-mouth movements appeared roughly sigmoidal and so were fit with an inverted Gaussian cumulative distribution function:

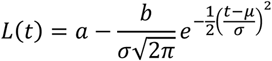

Transitions into the holding mode appeared to decay exponentially, so were fit with:

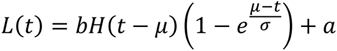

where *H*(*x*) is the Heaviside step function. In both cases, the parameters fit for each movement were the scale *b*, the offset *a*, the location *µ*, and the rate *σ*. Both models generally gave a good fit to the data (lowering-from-mouth: *R*^2^ = 95.8 ± 13.1%, median ± m.a.d., *n* = 192 movements from all videos; transport-to-mouth: *R*^2^ = 95.6 ± 8.2%, median ± m.a.d., *n* = 176 movements). For transport-to-mouth movements, the start and end of the movement were defined as the 80% confidence interval of the Gaussian fit (i.e., from 10% to 90% of the movement height below the baseline) and the amplitude was taken as the difference in the values of L at the edges of the 98% confidence interval. For lowering-from-mouth movements, the start was defined as *µ*, the end as the time of 99% decay, and the amplitude was the difference in the values L at those times. Mode durations were calculated from the time of the start of the movement transitioning into the mode to the time of the movement transitioning out. A new ethogram was constructed for each video using the start times extracted from the fitted models and this ethogram was used in all subsequent analyses.

Regrips were detected using the findpeaks function in the Matlab Signal Processing Toolbox as peaks in D with minimum prominence of 0.75 mm and a slope that exceeded 88 mm·s^-1^ in either direction (Barrett et al., 2020). The regrip peak height was defined as the difference between the value of D at the peak and the mean value during the pre- and post-regrip baselines, which were taken from 300 ms to 100 ms before and 100 ms to 300 ms after the regrip. The full-width at half maximum was defined as the width of the peak at halfway between baseline and the peak value. First regrip latency was defined as the time of the first regrip in a given oromanual event less the end time of the transport-to-mouth movement into said oromanual event. Peak parameters were extracted for individual regrips, before averaging first within and then across mice.

### Electrophysiological recordings and analysis

The linear arrays used were 32-channel silicon probes with ~1 MΩ impedance and 50-μm spacing (model A1×32-15mm-50-177-A32, NeuroNexus, Ann Arbor, MI), in linear configuration. Each probe was mounted on a linear translator (MTSA1, ThorLabs) that was in turn mounted on a 3-axis manipulator (MP285, Sutter, Novato, CA). Probes were positioned at the recording sites stereotactically based on the stereoscopically visualized location of bregma using the three axes of the Sutter manipulator, then slowly inserted into the cortex using the linear translator at a rate of 2 μm/s (controlled by software) to a nominal depth of 1,600 μm from the pia. Target coordinates for the three regions were as follows. Forelimb M1: 0.0 mm anterior-posterior (AP), 1.5 mm medial-lateral (ML) (Yamawaki et al., 2021); tongue/jaw M1: 1.8 AP, 2.5 ML (Mayrhofer et al., 2019); forelimb S1: 0.0 mm AP, 2.4 ML. For lateral recording sites (forelimb S1 and tongue/jaw M1), the probes were tilted by ~30° off the vertical axis for alignment with the radial axis of the cortex. For forelimb M1, the probes were inserted perpendicularly to the horizontal plane (for unilateral recordings) or ~15° off the vertical (for bilateral recordings, to avoid headstage collision). At the end of each experiment, the probes were removed, the craniotomy re-sealed with Kwik-Sil, and the mouse returned to its home cage.

Signals were amplified using RHD2132 headstages (Intan Technologies, Los Angeles, CA) and acquired at 30 kHz using an RHD2000 USB Interface Evaluation Board (Intan). Data was recorded using the Intan experimental interface evaluation software, triggered by the same trigger used to control video recording. To synchronize videos to electrophysiological recordings, the frame sync signal from the camera was recorded as a digital input to the RHD2000. RHD files recorded by the Intan software were converted to raw format using Matlab (The MathWorks, Natick, MA), from which spikes were detected and sorted using Kilosort (Pachitariu et al., 2016; Steinmetz et al., 2021). Results from Kilosort were manually verified using phy (https://github.com/cortex-lab/phy) as follows. Units with waveforms spanning more than 3 adjacent channels or with atypical waveform shapes were rejected as artifactual. Units displaying a clear refractory period (<1% of spikes within 1 ms) were classified as single units. All other units were classified as multiunits. Multiunits on the same channel with similar waveform shapes were merged. Single units were merged only if they were on the same channel, displayed similar shapes, and had no spikes in +/- 1 ms in their crosscorrelogram (Hall et al., 2021). Single units and multiunits (“active units”) were included in all analyses presented. Only data from probe recordings from which at least 10 active units could be isolated were included.

### Optogenetics

The laser-scanning photostimulation apparatus used for optogenetic silencing and stimulation experiments has been described in detail elsewhere (Jiang et al., 2018; Jiang et al., 2019). Briefly, 473 nm wavelength light from a blue laser source (LY473III-100, Aimlaser, Xi’an, China) was directed through an acousto-optic modulator (AOM, MTS110-A3-VIS, AA Opto-Electronic, Orsay, France) and an iris (SM1D12D, ThorLabs) before being deflected by a pair of galvanometer scan mirrors (GVSM002, ThorLabs) and focused onto the brain by a plano-convex spherical lens (LA1484-A, ThorLabs).

Laser power and location were controlled by sending voltage commands to the AOM and scan mirrors, respectively, from a NI USB-6229 data acquisition board controlled by WaveSurfer 0.945 (https://wavesurfer.janelia.org/). Bilateral forelimb M1 silencing was achieved by commanding the scan mirrors to rapidly direct the laser beam back and forth between right and left forelimb M1 for one second at 40 Hz, with the AOM command voltage set to zero while the mirrors were moving to avoid stimulating more medial areas (Morandell and Huber, 2017). For unilateral forelimb M1 corticospinal stimulation, the laser beam was directed to the expression site and pulsed at 40 Hz (20% duty cycle) for 250 ms (longer duration stimuli were used for one mouse, but the results were similar whether or not this mouse was included). The power at the focal plane was calibrated to 5-20 mW. For silencing and stimulation, stimuli were delivered for every ten seconds while the mice were eating, regardless of whether they were currently in holding or oromanual.

### Data analysis

All neural firing data was binned into 1 ms bins aligned to simultaneously recorded video frames prior to further analysis.

#### Analysis of event-aligned data

For Fig. 3–5, each active unit’s spike train was first aligned to the events of interest (transport-to-mouth movement starts, regrip peaks, lowering-from-mouth movement starts). Due to sparse firing of many active units, spike trains were further binned into 20 ms bins after alignment, thus providing smoother firing rate estimates while preserving the precise alignment to kinematics possible with kilohertz-rate video. Peri-event time histograms (PETHs) for each unit spanning 0.5s before to 1.0s after transport-to-mouth or lowering-from-mouth movements and 0.5s before to 0.5s after regrips were constructed by averaging the event-aligned binned firing rates over events. Significance of PETHs was assessed as follows. A bootstrap distribution of PETHs was constructed for each active unit and event type by randomly placing virtual transport-to-mouth or lowering-from-mouth onset times in each holding interval or oromanual event, respectively, or by placing a number of random regrip times equal to the number of actual regrips in each oromanual event. This was repeated 10,000 times for each unit and event type. Then, for each active unit and event type, two one-sided *p*-values for excitation and inhibition were calculated as the fraction of bootstrapped PETHs with maximum firing rates greater or equal to that of the real PETH or minimum firing rates less than or equal to that of the real PETH, respectively. Finally, an active unit was designated as significantly excited or inhibited by a given event type if the corresponding *p*-value was significant at a Holm-Bonferroni corrected alpha level of 0.05.

Peak excitation (inhibition) latencies were calculated for each active unit as the time bin of its PETH having the maximum (minimum) firing rate. The onset latency was calculated as the last time bin in which the PETH was less than (greater than) the minimum (maximum) pre-peak (pre-trough) firing rate of the PETH plus (minus) 10% of the difference between the minimum and maximum firing rates.

An oromanual phasic-tonic index (PTI) was calculated for each active unit as (FR_peak_ – FR_end_)/(FR_peak_+FR_end_), where FR_peak_ is the average firing rate in a 200 ms window surrounding the PETH bin with maximal firing rate and FR_end_ is the average firing rate in the 200 ms before the lowering-from-mouth movement (i.e., the last 200 ms of each oromanual event). This is similar to the definition used previously (Shalit et al., 2012), except that the window used to estimate peak firing is adapted on a per-unit basis, rather than using a fixed window.

Probe-average firing rates were estimated by averaging PETHs for all active units simultaneously recorded on the same probe. The probe-average PTI was calculated from this probe-average PETH exactly as for the active unit PTIs.

#### Non-negative matrix factorization analysis

Following Xu et al. (Xu et al., 2020), we used non-negative matrix factorization (NNMF) to perform simultaneous dimensionality reduction and clustering of unit responses. First, each unit’s firing rate was smoothed by convolution with a 20 ms causal boxcar, then aligned to transport-to-mouth, regrip, and lowering-from-mouth movements. Each unit’s mean response from −300 ms before to 300 ms after each event was calculated, temporally concatenated across conditions, and normalized to the interval [0, 1]. Units with zero mean event-aligned firing rate across all conditions were excluded as the normalized response is undefined in this case. The remaining normalized responses were concatenated into a single T × N matrix, where T is the number of time bins and N the number of units. This matrix was factorized into a W = T × k matrix of response templates and an H = k × N matrix of response weights using the nnmf function in Matlab. The optimal number of factors k was found based on 1000-fold leave-one-out bi-cross-validation (Owen and Perry, 2009), wherein one time bin and one unit are randomly left out of the input matrix to NNMF, W and H are recalculated, then the process is repeated 1000 times and k is chosen to minimize the mean (across folds) squared error between the full input matrix and the product of the factorizations calculated on the reduced input matrices. Units were pooled across mice and experiments for NNMF analysis. Each active unit was assigned to a cluster corresponding to the factor with the greatest contribution to that unit’s activity (i.e., the factor with the greatest weight in that unit’s column of the H matrix).

#### Kinematic reconstruction

To reconstruct forelimb trajectories from neural activity, we fit general linear models (GLM) from sliding windows of varying length and lag to the X, Y, and Z, coordinates of the third digit of each hand. The median nose position was taken as the origin and trajectories for each experiment were rotated to align the X, Y, and Z world axes with vectors perpendicular to the mouse’s sagittal, coronal, and horizontal planes (i.e., movements along the X, Y, and Z axes corresponded to side-to-side, back-and-forth, and up-and-down movements, respectively). Trajectories and firing activity were binned into 20 ms bins, as for the PETH analyses. GLMs were fit using the fitrlinear function in Matlab using ridge regularization. Half the data was used to choose the ridge parameter, *λ*, using Bayesian hyperparameter optimization with ten cross-validation folds. Having found the optimal *λ*, the GLMs were re-fit to the remaining half of the data using ten-fold cross-validation. Reported *R*^2^ and regression coefficient values in the text are means over folds.

### Histology

Probes were coated with fluorescent dye (DiO, DiI, or DiD, Vybrant Multicolor Cell Labeling Kit, Invitrogen, Carlsbad, CA) for subsequent histological verification of probe placement under epifluorescence microscopy. Following the final recording with each mouse, the mouse was sacrificed by overdose of isoflurane followed by decapitation, and the brain was quickly dissected and placed in 4% paraformaldehyde solution (Electron Microscopy Sciences, Hatfield, PA) in 0.01 M phosphate-buffered saline (PBS, Sigma-Aldrich, Burlington, MA). Brains were fixed overnight, then washed the next day with PBS and stored in PBS with 0.02% w/v sodium azide (DOT Scientific, Burton, MI) until imaging. To image electrode tracks, brains were sliced into 100 μm serial sections using a microtome (Microm HM 650 V, Microm International, Walldorf, Germany) and imaged with a Retiga 2000R CCD camera (QImaging, Burnaby, BC, Canada) mounted on an Olympus SZX16 upright microscope (Olympus, Tokyo, Japan). Slices were imaged under bright-field illumination and appropriate fluorescent illumination for the dye(s) used in each experiment. If the dyed probe track could not be found, probe location was based on appearance of probe tracks in the bright-field image. Stereotactic coordinates were calculated for the top and bottom of the probe track as follows. The AP coordinate relative to bregma was found by counting slices from the slice containing the middle of the anterior commissure. The ML coordinate was taken as the horizontal distance to the midline in pixel values, converted to micrometers using a previously calibrated conversion factor for the magnification at which the image was taken. Finally, the coordinates of each recording were defined as the mean of the AP and ML coordinates of the top and bottom of the corresponding probe track. In one case, because probe tracks could not be recovered histologically, areal assignment was based on the coordinates of the area targeted.

### Statistical analyses

All data presented in figures and the main text are means ± standard deviation unless otherwise indicated. Recordings were made on multiple days for some mice, with the probes being repositioned each day. In these instances, kinematic data from the same mouse are pooled across days, but data relating to neural activity from the same cortical area are averaged first within and then across days. For hypothesis testing, nonparametric tests were used wherever possible. For paired samples, Wilcoxon’s signed rank was used unless the number of samples was below the minimum required for significance at a *p* < 0.05 level (i.e., *n* < 6), in which case paired *t*-tests were used provided the distribution of paired differences was not significantly different from normal as assessed by the Shapiro-Wilk test. For comparing medians among more than two groups, Kruskal-Wallis ANOVA was used. For comparing means among groups across two factors simultaneously, non-parametric two-way ANOVA was performed using permutation tests via the aovp function in R. For comparing parameters measured at multiple timepoints (optogenetic silencing and stimulation experiments) or across multiple conditions (side and coordinate for reconstruction) in the same mice, repeated-measures ANOVA in Matlab (for one within-subject factor) or R (for multiple within-subject factors) was used after checking normality of data within each level of each within-subjects factor using the Shapiro-Wilk test and sphericity using Mauchly’s test. Where sphericity was violated, the Greenhouse-Geisser correction was applied to the degrees of freedom of the corresponding *F*-test. In all cases the Bonferroni method was used for follow-up comparisons. For regression where the slope was expected to vary per mouse, linear mixed-effects regression models were used with mouse as a grouping variable and residuals visually inspected for normality to ensure goodness of fit.

## ACKNOWLEDGEMENTS

For technical assistance we thank F. Hausmann, M. Martin, M. Raineri Tapies, and M. Torres. For comments and advice on a draft of the paper, we thank L. Miller, A. Miri, L. Pinto, and M. Tresch.

Grant support: National Institute of Neurological Disorders and Stroke of the National Institutes of Health under Award Numbers R01NS061963 and R21NS116886.

